# Slow drift of neural activity as a signature of impulsivity in macaque visual and prefrontal cortex

**DOI:** 10.1101/2020.01.10.902403

**Authors:** Benjamin R. Cowley, Adam C. Snyder, Katerina Acar, Ryan C. Williamson, Byron M. Yu, Matthew A. Smith

**Affiliations:** Princeton Neuroscience Institute, Princeton University, Princeton, NJ; Department of Machine Learning, Carnegie Mellon University, Pittsburgh, PA; Center for Neural Basis of Cognition, Carnegie Mellon University, Pittsburgh, PA; Carnegie Mellon Neuroscience Institute, Carnegie Mellon University, Pittsburgh, PA; Department of Electrical and Computer Engineering, Carnegie Mellon University, Pittsburgh, PA; Department of Biomedical Engineering, Carnegie Mellon University, Pittsburgh, PA; Department of Brain and Cognitive Sciences, University of Rochester, Rochester, NY; Center for Neuroscience, University of Pittsburgh, Pittsburgh, PA; University of Pittsburgh School of Medicine, University of Pittsburgh, Pittsburgh, PA; Department of Ophthalmology, University of Pittsburgh, Pittsburgh, PA

## Abstract

An animal’s decision depends not only on incoming sensory evidence but also on its fluctuating internal state. This internal state is a product of cognitive factors, such as fatigue, motivation, and arousal, but it is unclear how these factors influence the neural processes that encode the sensory stimulus and form a decision. We discovered that, over the timescale of tens of minutes during a perceptual decision-making task, animals slowly shifted their likelihood of reporting stimulus changes. They did this unprompted by task conditions. We recorded neural population activity from visual area V4 as well as prefrontal cortex, and found that the activity of both areas slowly drifted together with the behavioral fluctuations. We reasoned that such slow fluctuations in behavior could either be due to slow changes in how the sensory stimulus is processed or due to a process that acts independently of sensory processing. By analyzing the recorded activity in conjunction with models of perceptual decision-making, we found evidence for the slow drift in neural activity acting as an impulsivity signal, overriding sensory evidence to dictate the final decision. Overall, this work uncovers an internal state embedded in the population activity across multiple brain areas, hidden from typical trial-averaged analyses and revealed only when considering the passage of time within each experimental session. Knowledge of this cognitive factor was critical in elucidating how sensory signals and the internal state together contribute to the decision-making process.

## Introduction

Over the course of a day, we face many perceptual decisions (e.g., a driver waiting for a traffic signal to turn green). These decisions are influenced not only by information at hand (e.g., the color of the traffic signal) but also by a myriad of internal factors that influence our choices at the moment a decision is made (e.g., fatigue from driving). These factors have timescales ranging from many minutes (e.g., arousal) to seconds (e.g., spatial attention) to fractions of a second (e.g., recent visual stimulus history), and together define an “internal state”. When an animal is tasked with making back-to-back perceptual decisions in a laboratory setting, fluctuations in the internal state still influence perceptual outcomes despite the constancy of the statistics of task variables. Indeed, many studies have investigated how and to what extent internal states influence the outcome of a decision [31, 37, 64], as this provides insight into the cognitive and neural mechanisms underlying decision-making [90].

Perceptual decisions have often been characterized with models that accumulate (noisy) sensory evidence, such as variants of the drift-diffusion model [50, 11, 79], and there are neural signatures of this accumulation process [31, 47, 29, 36, 10, 35, 39]. Decisions can be better characterized by extending these models to include additional factors, such as an urgency signal [15, 34, 68], duration of fixation [51], the number of alternative choices [14], the reward or choice history of previous trials [1, 102], or a model term describing errors (i.e., ‘lapses’) that occur independently of sensory evidence [105, 76]. Less is known about the neural signatures of these cognitive factors [75]. Most of these models focus solely on short-term influences on decisions (e.g., within a single trial or between consecutive trials), despite the presence of long-term influences (i.e., over several minutes to hours) on decisions, such as fatigue [58], arousal [4, 64, 101], and satiety [2], among others. This motivates the question of how such long-term changes in a subject’s internal state influence the subject’s choices in a perceptual decision-making task.

In this work, we trained macaque monkeys to make hundreds of perceptual decisions over the course of several hours. We found that the animals’ behavior changed slowly during the task, unprompted by task structure. These slowly changing behaviors included their tendency to report a change when none had happened (i.e., the rate of false alarms), reaction time, and pupil diameter. These behavioral measures covaried with each other over the course of tens of minutes, indicating they have a shared neural origin. During the task, we simultaneously recorded neural population activity from visual area V4 and prefrontal cortex (PFC), two brain areas that have been implicated in forming perceptual decisions. We identified a slow fluctuation (termed the *slow drift*) in V4 activity that covaried with the slow changes in behavior. Surprisingly, we also found that PFC neurons, despite their physical distance from V4, exhibited a slow drift that was highly correlated with that of V4. We uncovered evidence that this slow drift acts as an arousal or impulsivity signal, which influenced the final decision through a pathway independent of sensory evidence. In addition, we found evidence that downstream areas remove or account for this slow drift to prevent it from adversely affecting the sensory readout. This may explain how perceptual decisions can be formed reliably when sensory signals (e.g., V4 activity) drift so profoundly. Overall, this work identifies a slowly varying internal signal present in both V4 and PFC (and likely throughout the cortex), and proposes a role for this signal in the decision-making process.

## Results

We trained two adult, male rhesus monkeys (*Macaca mulatta*) to perform an orientation-change detection task (Fig. 1**a**). Results from a portion of data collected from this experiment have been reported previously [97]. After an initial flash of two Gabor stimuli, each succeeding flash had the same probability (30%, 40% for monkeys 1, 2) that the orientation of one of the stimuli would change (i.e., a flat hazard function). The animal was trained to detect a change in the stimulus and saccade to it. A ‘hit’ trial was one in which the animal correctly made a saccade to the changed stimulus and then received a reward. A ‘false alarm’ trial was one in which the animal incorrectly made a saccade to a stimulus when no change occurred. To measure the animal’s ability to detect a stimulus change, we used signal detection theory to calculate sensitivity of detection *d*^’^ [56, 54]. As expected, the animal’s sensitivity increased as the change in orientation increased (Fig. 1**b**, ‘cued stimulus changed’). To probe the effects of spatial attention, we structured trials into alternating blocks such that for each block a changed stimulus was 90% likely to occur at one location (Fig. 1**a**, ‘cued stimulus’) and 10% likely to occur at the other location (i.e., the ‘uncued stimulus’). Animals were more sensitive (based on *d*^’^) at detecting stimulus changes at the cued location than changes that occurred at the uncued location (Fig 1**b**, similar results for second monkey in Supplementary Fig. 1), suggesting that the animals deployed spatial attention to the cued location.

**Figure 1:**
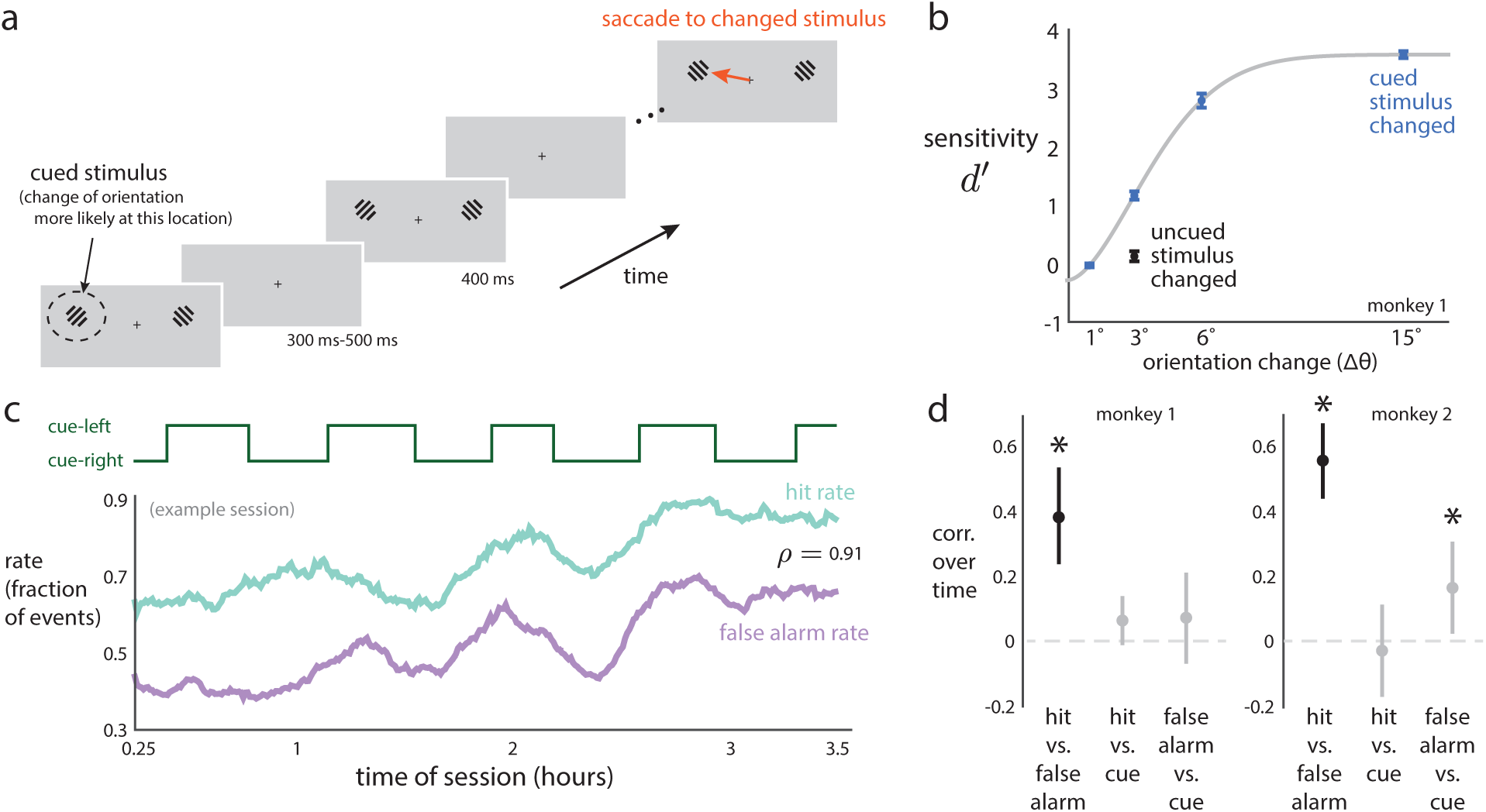
Behavior slowly fluctuates during an experimental session. **a**. Orientation-change detection task with cued attention. On each stimulus flash, either of two oriented gratings could change orientation. The animal’s task was to saccade to the stimulus location whose orientation changed (orange arrow). The animal was previously cued as to which of the two stimulus locations was more likely to change its orientation (cued stimulus location is indicated by dashed circle, which was not shown to the animal). **b**. The animal’s sensitivity *d*^’^ increased with larger orientation changes of the cued stimulus (blue dots, gray line indicates fit with Weibull function). For trials in which the uncued stimulus changed, the animal’s sensitivity was lower (Δ*θ* = 3°, black dot below blue dot, *p <* 0.002 for both monkeys, paired permutation test). Dots indicate means over sessions, error bars indicate ±1 s.e.m. **c**. A representative session in which the hit rate (teal line) and false alarm rate (purple line) slowly covaried over time (Pearson’s *ρ* = 0.91, *p <* 0.002, permutation test). For this session, hit rate (*ρ* = 0.10, *p* = 0.096) and false alarm rate (*ρ* = 0.23, *p <* 0.002) rates had little to no relationship with which stimulus location was cued (top, green line). **d**. Correlations over time between hit rate, false alarm rate, and cued stimulus location (asterisks denote significance over chance levels, *p <* 0.05, permutation test). Dots indicate medians over sessions, error bars indicate bootstrapped 90% confidence intervals.

### Slow fluctuations in behavioral variables reveal the presence of a shifting internal state

Although task statistics remained constant throughout each recording session (average session duration: 2.3, 3.0 hours for monkeys 1, 2), the animal’s internal state was free to vary with the passage of time due to such factors as satiety, fatigue, motivation, etc. Indeed, when we analyzed the animal’s behavior across the entire session, we found fluctuations in its behavior (Fig. 1**b**) that were hidden from our previous analysis for which we collapsed across trials for the purpose of measuring the effect of spatial attention. The time course of two commonly-analyzed behavioral variables—hit rate and false alarm rate—slowly fluctuated together over the entire session (Fig. 1**c**, example session, teal and purple lines). The difficulty of the perceptual decisions combined with uncertainty of when a change would occur yielded high false alarm rates (∼50% for this session). Over the course of the session, a change to a more impulsive behavioral state would result in more correct choices on the difficult trials, but also more false alarms. Conversely, a more hesitant approach would yield a lower false alarm rate, but also reduce the likelihood of correctly detecting difficult stimulus discriminations. For all sessions, we found that hit rate and false alarm rate covaried (Fig. 1**d**, black dots), indicating a fluctuation between a more impulsive and more hesitant state. These two variables drifted together on long timescales (hit rate: 13.4 ± 1.3 minutes, 15.9 ± 2.2 minutes for monkeys 1, 2; false alarm rate: 14.0 ± 1.3 minutes, 14.5 ± 1.8 minutes for monkeys 1, 2; mean ±1 s.e.m. over sessions; see Methods).

One possible explanation for these behavioral fluctuations was that spatial attention was cued to each of the two stimuli in blocks of trials (Fig. 1**c**, green line, each block lasted on average 20.4, 23.7 minutes for monkeys 1, 2), and the animal reacted to the switches between blocks. This cueing occurred at the beginning of each block (see Methods), and resulted in robust attentional effects in both behavior and neural activity [97]. While the blocks of cued trials could in principle have explained the slow fluctuations in the animal’s behavior that we report here, we found this not to be the case (Fig. 1**c**, example session, teal and purple lines have little to no covariation with green line, Fig. 1**d**, across all sessions, gray dots). We also found these behavioral shifts did not simply reflect the animal becoming gradually fatigued over the course of the session and “guessing” more at the end, as it was not the case that hit rate and false alarm rate strictly increased during each session (Supplementary Fig. 2). Instead, because task statistics remained constant throughout the session, these behavioral shifts likely reflected a slow change in the animal’s internal state from minute to minute within the session. Such fluctuations could lead the animal to change its behavior—at different times having a higher or lower impetus to make a choice (or conversely, a higher or lower resistance to initiating a movement), leading to correlated fluctuations in hit rate and false alarm rate.

Next, we asked whether these slow fluctuations in behavior were large relative to the prominent behavioral effect of spatial attention that we observed during our task. The behavioral shifts of spatial attention involved large changes in hit rate (0.35 ± 0.09, 0.41 ± 0.12 for monkeys 1, 2; see Methods) but small changes in false alarm rate (0.05 ± 0.04, 0.03 ± 0.02) between trials in which the changed stimulus occurred in the cued location versus the uncued location. In contrast, the slow fluctuations in behavior had large changes in both hit rate (0.32 ± 0.12, 0.34 ± 0.10 for monkeys 1, 2) and false alarm rate (0.21 ± 0.09, 0.32 ± 0.09). These results suggest that the animal’s behavior slowly changed across the session on the order of tens of minutes, at a level that was on par with (but uncorrelated with) the behavioral effects we measured due to spatial attention.

### The activity of V4 neurons slowly drift together

We next considered whether a neural signature of this fluctuating internal state might be found in the activity of cortical neurons (Fig. 2**a**). We investigated two brain regions, visual area V4 and dorsolateral prefrontal cortex (PFC). We selected these areas because past work has suggested that perceptual decisions might be influenced by noise in sensory neurons [92] and by top-down control mechanisms [22]. In each animal, we implanted two 100-electrode “Utah” arrays in the same hemisphere to simultaneously record the activity of populations of neurons in V4 and PFC. We began by asking if V4 activity contained a neural signature of the slow behavioral fluctuations (Fig. 1**c**).

**Figure 2:**
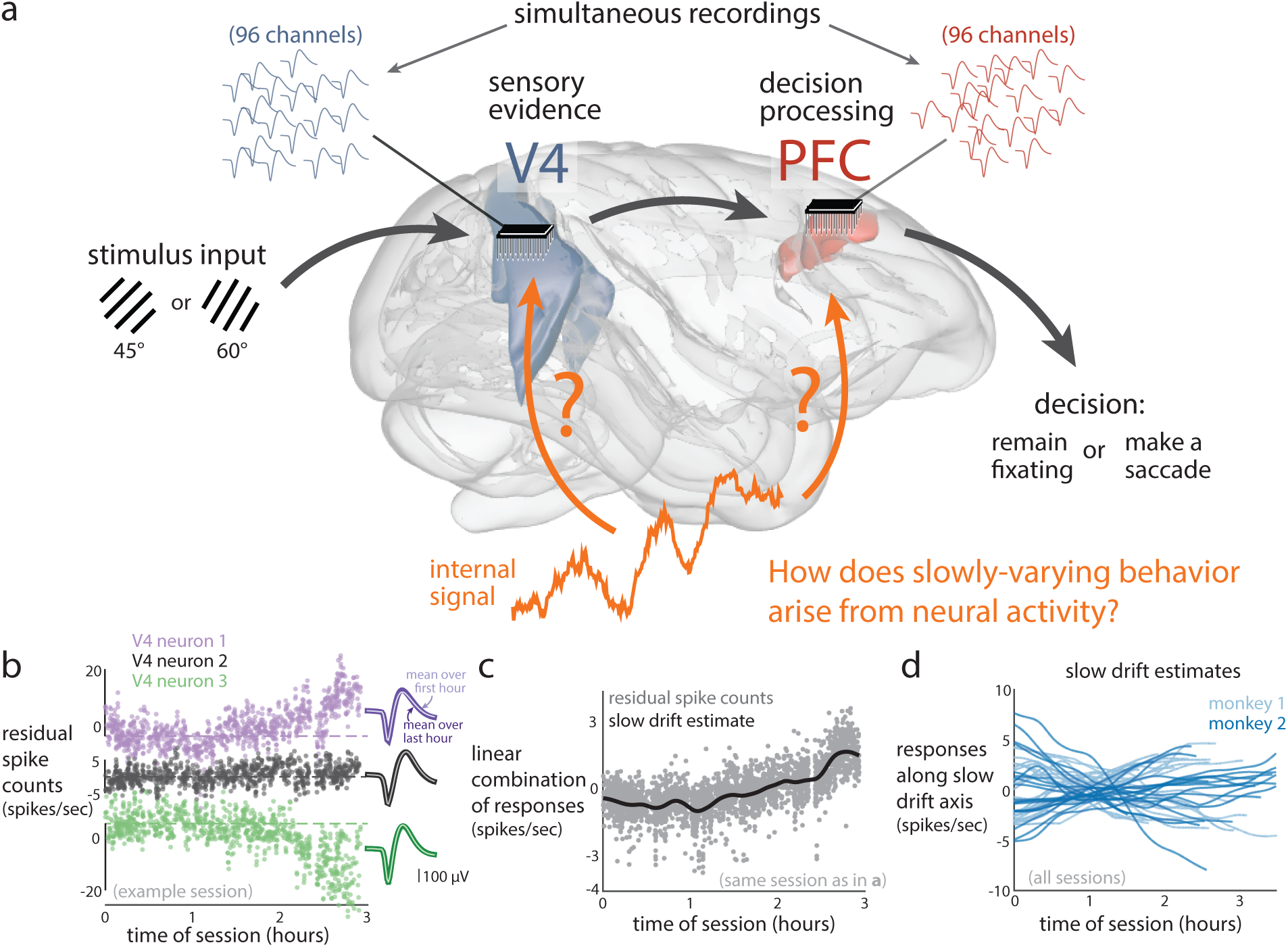
Neural activity slowly drifts throughout a session. **a**. We simultaneously recorded population activity from visual area V4, thought to transform stimulus input into sensory evidence, and from prefrontal cortex (PFC), thought to be involved in cognitive control signals and decision-making. We asked whether these brain areas exhibit a neural signature (orange) of the observed slow fluctuations in behavior (Fig. 1c). **b**. Time course of the activity of three neurons from a representative session. Each dot is the 400 ms-binned residual spike count (raw spike count minus mean spike count averaged over stimulus repeats) for one stimulus flash. Dashed lines denote mean residual spike counts for the first 30 minutes. Insets: Spike waveforms corresponding to the three neurons. Each waveform is the mean spike waveform averaged across either the first hour of the session (lighter shade, ‘first hour’) or the third hour of the session (darker shade, ‘last hour’). Lines are close to overlapping. **c**. Linear combination of the activity of the 48-simultaneously-recorded neurons from the same session as in **b**. Each gray dot is a linear combination (identified using PCA, see Methods) of the residual spike counts for one stimulus flash. These projections were then Gaussian-smoothed to estimate the slow drift (black line). **d**. Time courses of the slow drifts (computed in the same manner as in **c**) for all sessions for both monkeys.

First, we considered the activity of individual neurons. We found that some neurons increased their activity (Fig. 2**b**, ‘neuron 1’), some neurons decreased their activity (Fig. 2**b**, ‘neuron 3’), and other neurons did not exhibit drift in their activity (Fig. 2**b**, ‘neuron 2’) during a typical session. We ensured that this was not due to instability in neural recordings (see Fig. 2**b** spike waveform insets, Supplementary Fig. 3, and Methods). In addition, we confirmed that the basic properties of the recorded V4 neurons (i.e., mean firing rates, Fano factors, and noise correlations) were consistent with previous studies (Supplementary Fig. 4, as well as analysis published in [97]).

To quantify the coordinated drift in activity across the population of simultaneously-recorded neurons, we applied principal component analysis (PCA) to binned spike counts (after subtracting each neuron’s mean response to the stimulus, see Methods), and found a dominant linear combination of the neurons for which their activity drifted strongly. The weights (also known as loadings) of this linear combination represent an axis in the population activity space, which we define as the *slow drift axis*. We then projected the activity onto the slow drift axis (Fig. 2**c**, gray dots) to reveal a substantial fluctuation in neural activity over the course of the 3 hour experimental session (Fig. 2**c**, black line). We define this fluctuation as a *slow drift*.

The time course of slow drift varied across sessions (Fig. 2**d**), with a timescale of around 40 minutes (Supplementary Fig. 5, 32.8 ± 3.1, 43.5 ± 2.7 minutes for monkeys 1, 2; see Methods). The slow drift was different from the average of activity taken across neurons (Supplementary Fig. 6), which has been proposed as a way to summarize population activity using a single variable ([82, 37, 73]; but see [97]). We analyzed the weights of the linear combination (i.e., the slow drift axis), and found that about 50% of neurons had activity that drifted together in the same direction, about 25% drifted in the opposite direction, and the remaining 25% had little to no slow drift (Supplementary Fig. 6). Thus, the slow drift would be difficult to detect using individual neurons recorded sequentially or the average of simultaneously recorded neurons, because the slow drift is a coordinated but diverse fluctuation across the population of neurons.

### The slow drift covaries with slow fluctuations in behavior

We found that the slow drift is a prominent neural fluctuation with a similar timescale (tens of minutes) as that of fluctuations in behavior, as well as tightly coupled with both hit rate and false alarm rate (Fig. 3**a**). We sought to link our neural and behavioral observations of slow drift more directly. We quantified this relationship in two ways. First, within each session, we found that fluctuations in the slow drift and multiple behavioral variables, including hit rate and false alarm rate, were correlated over time within a session (Fig. 3**b**, see also Supplementary Fig. 7 for individual animals). That the slow drift was correlated with all five behavioral variables tested suggested that the slow drift was correlated with a behavioral pattern in which hit rate, false alarm rate, and pupil diameter increase while trial duration and reaction time decrease (or vice versa). Indeed, we found that the slow drift was correlated with this pattern, which explained 60% of the behavioral variance (Supplementary Fig. 8). This pattern of correlations among behavioral variables was consistent with an underlying shift between an impulsive behavioral state, associated with making more saccades with short reaction times while the pupil is dilated, versus a hesitant behavioral state, in which there are less saccades, longer reaction times, and a more constricted pupil.

**Figure 3:**
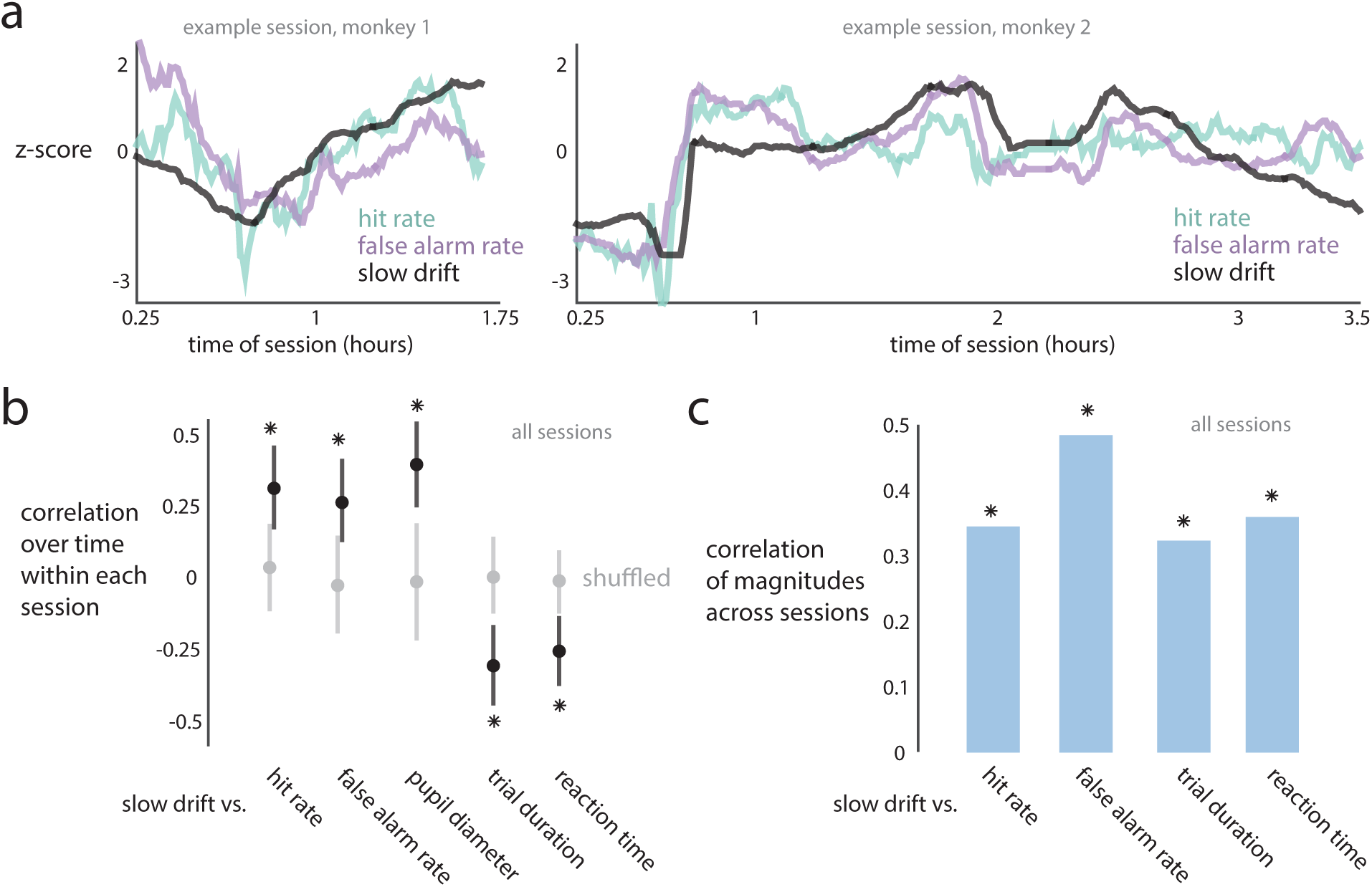
The slow drift of V4 neural activity covaries with slow fluctuations in behavior. **a**. Two example sessions for monkey 1 (left panel) and monkey 2 (right panel) in which hit rate, false alarm rate, and the slow drift covaried together (between hit rate and false alarm rate: *ρ* = 0.63, 0.84 for monkeys 1, 2; between slow drift and hit rate: *ρ* = 0.72, 0.64; between slow drift and false alarm rate: *ρ* = 0.27, 0.67). **b**. Within each session, the slow drift and behavioral variables were correlated over time (asterisks correspond to a significance of *p <* 0.002 over shuffling across sessions, permutation test). Dots indicate medians over sessions, error bars indicate bootstrapped 90% confidence intervals. **c**. Across sessions, correlations between the magnitudes of fluctuations of the slow drift and behavioral variables. Slow drift magnitude was the within-session variance over time of the slow drift. Behavioral variable magnitude was the within-session variance over time of that variable. All correlations were significant (asterisks correspond to a significance of *p <* 0.05 over chance levels, permutation test). The magnitude of pupil diameter was not included because pupil diameter measurements were not comparable across sessions (see Methods).

Second, across sessions, we found that the magnitude of firing rate changes in the slow drift (measured as the variance of the slow drift, see Methods) covaried with the magnitude of behavioral changes (Fig. 3**c**, see Supplementary Fig. 7 for individual animals). Thus, on sessions where the slow drift substantially fluctuated, there were also substantial behavioral changes. Taking both findings together, the slow drift appears to reflect the fluctuations of an underlying internal state.

### The slow drift is unrelated to and larger than the neural effect of spatial attention

To evaluate the origin of this neural effect, we asked to what extent the slow drift relates to another well-known modulation of the internal state of the animal, spatial attention [83, 60, 66], which we cued to different locations in blocks of trials (Fig. 4**a**). We refer to trials for which the stimulus location was cued either inside or outside the V4 receptive fields as ‘cue-in’ or ‘cue-out’ trials, respectively (Fig. 4**a**, orange and green lines). Similar to how the slow fluctuations in behavior were not correlated with the cued stimulus location (Fig. 1**c** and **d**), we found that the slow drift in neural activity did not covary with the timing of the cued blocks (e.g., Fig. 4**a**, black lines versus orange and green lines; mean *|ρ|* = 0.18, 0.15 for monkeys 1, 2, both no greater than expected by chance, *p* = 0.71 and *p* = 0.56, one-sided permutation test). Thus, the slow drift was not a neural signature of spatial attention.

**Figure 4:**
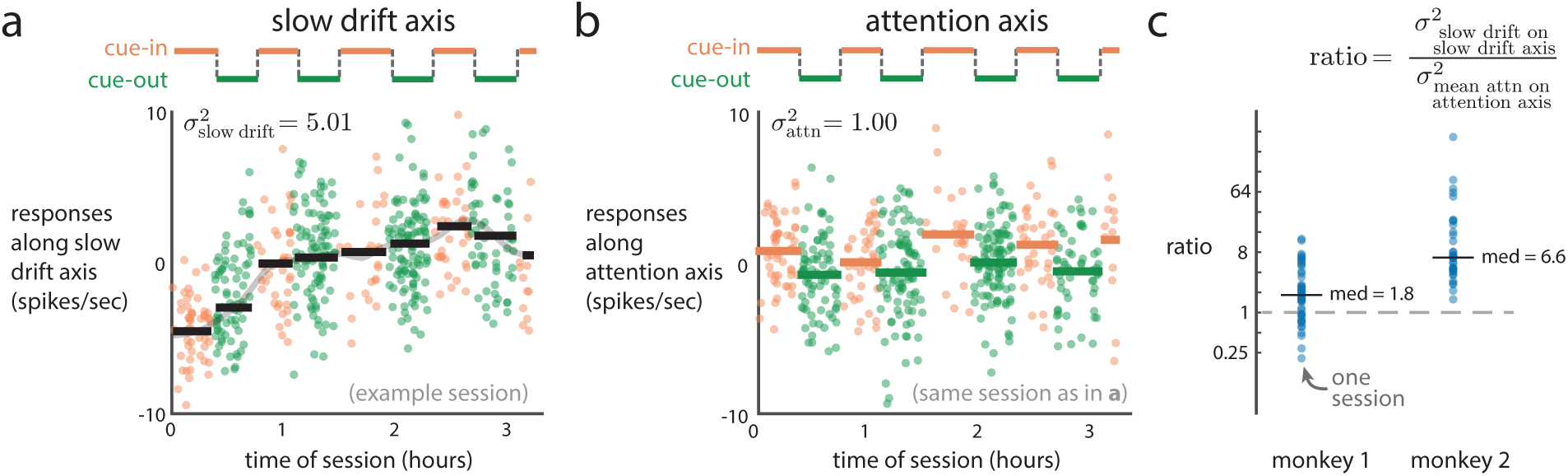
The slow drift in V4 activity is larger than the effect of spatial attention. **a**. Neural responses (orange and green dots) along the slow drift axis were unrelated to how blocks of trials were cued (black lines versus orange and green lines, *|ρ|* = 0.06) in an example session. Dot color indicates whether the cued stimulus location was within (‘cue-in’, orange) or outside (‘cue-out’, green) the receptive fields of the V4 neurons. Black lines indicate the mean activity along the slow drift axis within each cued block. The magnitude of firing rate changes of the slow drift 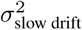 was measured as the variance of the vertical levels of the black lines. **b**. For the same session as in **a**, responses along an attention axis capturing the largest difference between mean spike counts during cue-in versus cue-out trials. The effect of attention 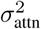 was measured as the variance of the mean value of responses averaged within each block, where the variance was taken across blocks (i.e., the spread of the orange and green lines). **c**. Ratios 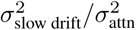 across sessions. Medians (‘med’) are also shown.

Instead, we used spatial attention to interpret the size of the firing rate modulations corresponding to the slow drift. We measured the magnitude of firing rate changes as the variance of the slow drift over time (Fig. 4**a**, spread of black lines, 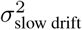), and compared it to the variance of changes in responses across cue-in and cue-out blocks (Fig. 4**b**, spread of orange and green lines, 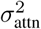). Because the slow drift was constrained to be along one axis in firing rate space (i.e., the slow drift axis), we also constrained the response changes due to attention to be along an axis for which the responses most differed between cue-in and cue-out blocks (i.e., the *attention axis*, see Methods). For this example session, the size of the slow drift was 5 times larger than the effect size of attention (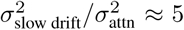). The ratio of the size of the slow drift divided by the effect size of attention was greater than 1 for both monkeys (Fig. 4**c**, median ratios: 1.8, 6.6 for monkeys 1, 2, *p <* 0.002, paired permutation test). These results indicate that the slow drift is a prominent neural fluctuation that leads to larger variation than that of the neural fluctuations due to spatial attention.

### V4 and PFC neurons share the same slow drift

The covariation between the slow drift in V4 activity and behavioral variables (Fig. 3) suggests that the slow drift could play a role in the decision-making process. However, it is unclear how the slow drift propagates through the neural circuit to ultimately influence the decision outcome. To better understand this, we recorded neural activity in PFC, a brain area relevant to integrating sensory evidence and forming decisions [23, 57, 78], simultaneous with the V4 recordings (Fig. 2**a**). One possibility is that PFC has no slow drift, suggesting that the slow drift is a local signal within the neural circuitry for visual processing. Another possibility is that PFC has a slow drift but this slow drift does not covary with the V4 slow drift, indicating that multiple local signals co-exist within the brain. Finally, it could be that PFC has a slow drift that covaries with the V4 slow drift, suggesting that the slow drift has a global presence in the brain.

To test these possibilities, we applied the same analyses to the PFC activity as we did to the V4 activity (i.e., PCA was applied separately to V4 and PFC activity, see Methods). Visually, we found that PFC activity had a slow drift that was remarkably similar to the V4 slow drift on a session-by-session basis (Fig. 5**a**, black and red lines). We quantified the relationship between the V4 and PFC slow drifts in two ways. First, within each session, we found that the V4 slow drift strongly covaried with the PFC slow drift over time (Fig. 5**b**, orange histogram, median *|ρ|* = 0.96, 0.93 for monkeys 1, 2, *p <* 0.002, permutation test). These correlations were significantly greater than those expected if PFC had no slow drift, computed by shuffling PFC responses within each session and reidentifying the slow drift (Fig. 5**b**, gray histogram, median *|ρ|* = 0.50, significantly less than the median of the real data, *p <* 0.002, permutation test). We also considered a more conservative null hypothesis in which PFC had a slow drift that was uncorrelated with the V4 slow drift. We obtained this null distribution by shuffling the PFC activity across sessions (but not within a session). We found that the correlation in slow drift between V4 and PFC was still significantly higher than this chance level (*p <* 0.002, permutation test; see Methods).

**Figure 5:**
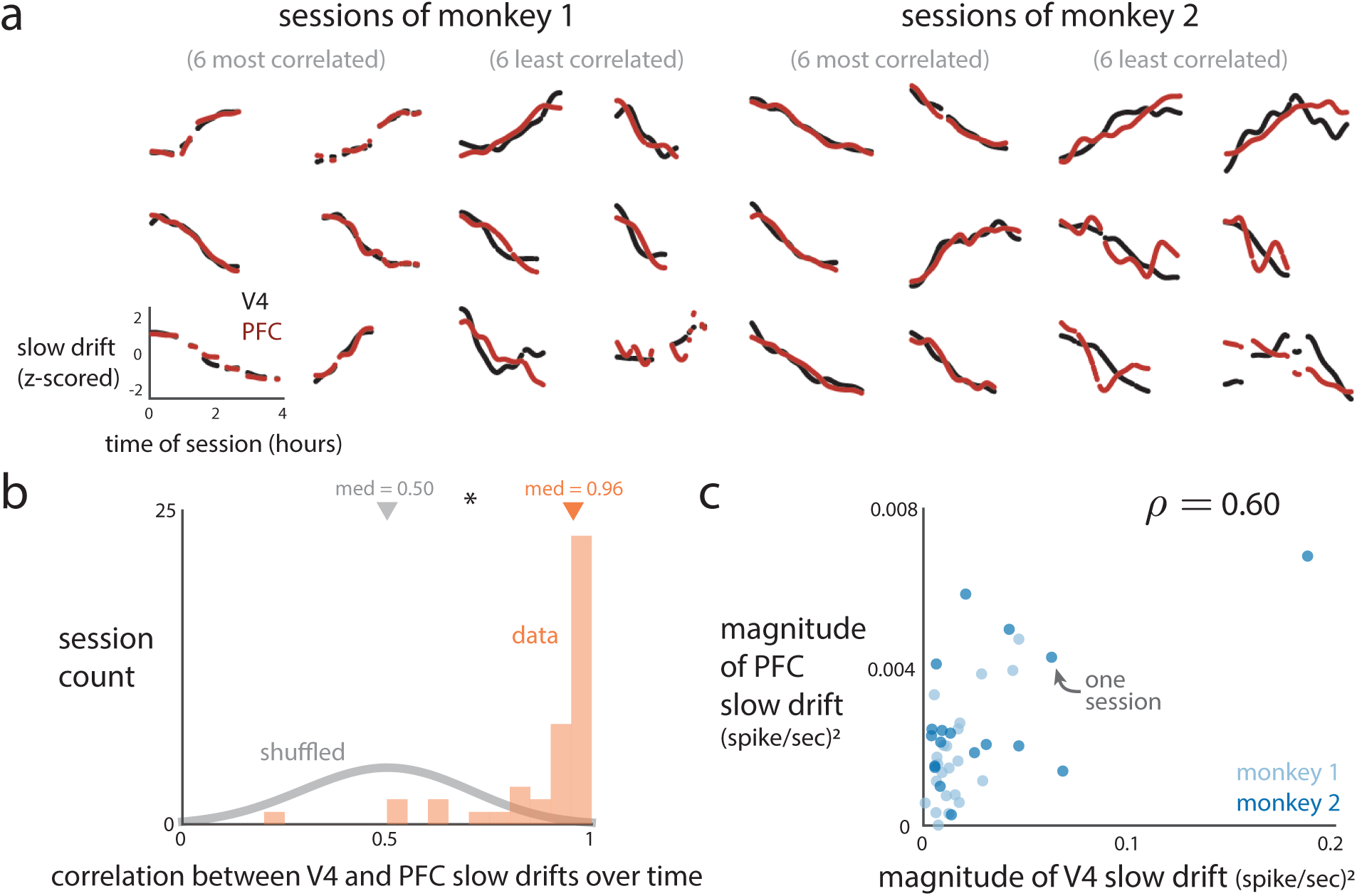
V4 and PFC activity share a similar slow drift. **a**. The slow drift of PFC neurons (red) and slow drift of V4 neurons (black) for the 6 most correlated and 6 least correlated sessions for each monkey. **b**. Within each session, the V4 and PFC slow drifts were more correlated over time than if PFC activity was shuffled across time (asterisk denotes *p <* 0.002, permutation test). **c**. Across sessions, the magnitude of fluctuations of the V4 slow drift was correlated with that of the PFC slow drift (*ρ* = 0.60, *p <* 0.002, permutation test). Magnitude was measured as the variance of the slow drift over time within a session. Each dot represents one session. The correlation was not solely due to the outlying session (magnitude of V4 slow drift *≈* 0.2), as its removal still led to a significant correlation (*ρ* = 0.43, *p <* 0.005, permutation test).

Second, across sessions, we found that the magnitude of fluctuations of the V4 slow drift correlated with that of the PFC slow drift (Fig. 5**c**, *ρ* = 0.68, 0.60 for monkeys 1, 2). Magnitude was measured for each brain area separately as the variance of the slow drift within a session (same metric as in Fig. 3**c**). Thus, when the V4 slow drift had large fluctuations, the PFC slow drift also tended to have large fluctuations for the same session. This finding that PFC had a slow drift which closely matched that in V4 suggests that the slow drift has a global presence in the brain.

### The influence of the slow drift on the decision-making process

In previous sections, we identified a slow drift in V4 and PFC activity that was related to slow behavioral fluctuations. These results shed light on the possible roles the slow drift can play in the neural circuit governing decision making. One possibility is that the slow drift reflects a signal that biases sensory evidence to be closer to or farther from a decision threshold. For example, an increase in the slow drift could bias evidence closer to this threshold, leading to an increase in both hit rate and false alarm rate (i.e., a change in criterion). This is consistent with our observations that hit rate, false alarm rate, and the slow drift covaried together (Fig. 3**b**). This possibility can also explain the presence of the slow drift in both V4 and PFC: the slow drift biases sensory evidence (e.g., V4 activity), which is then read out by downstream areas that help to form the decision (e.g., PFC) (Fig. 6**a**). The slow drift could arise in sensory areas from bottom-up feedforward sensory noise (Fig. 6**a**, right inset, 1, from the sensory periphery) or from top-down feedback signals (Fig. 6**a**, right inset, 2, from downstream areas), both of which are thought to induce choice probabilities [22].

**Figure 6:**
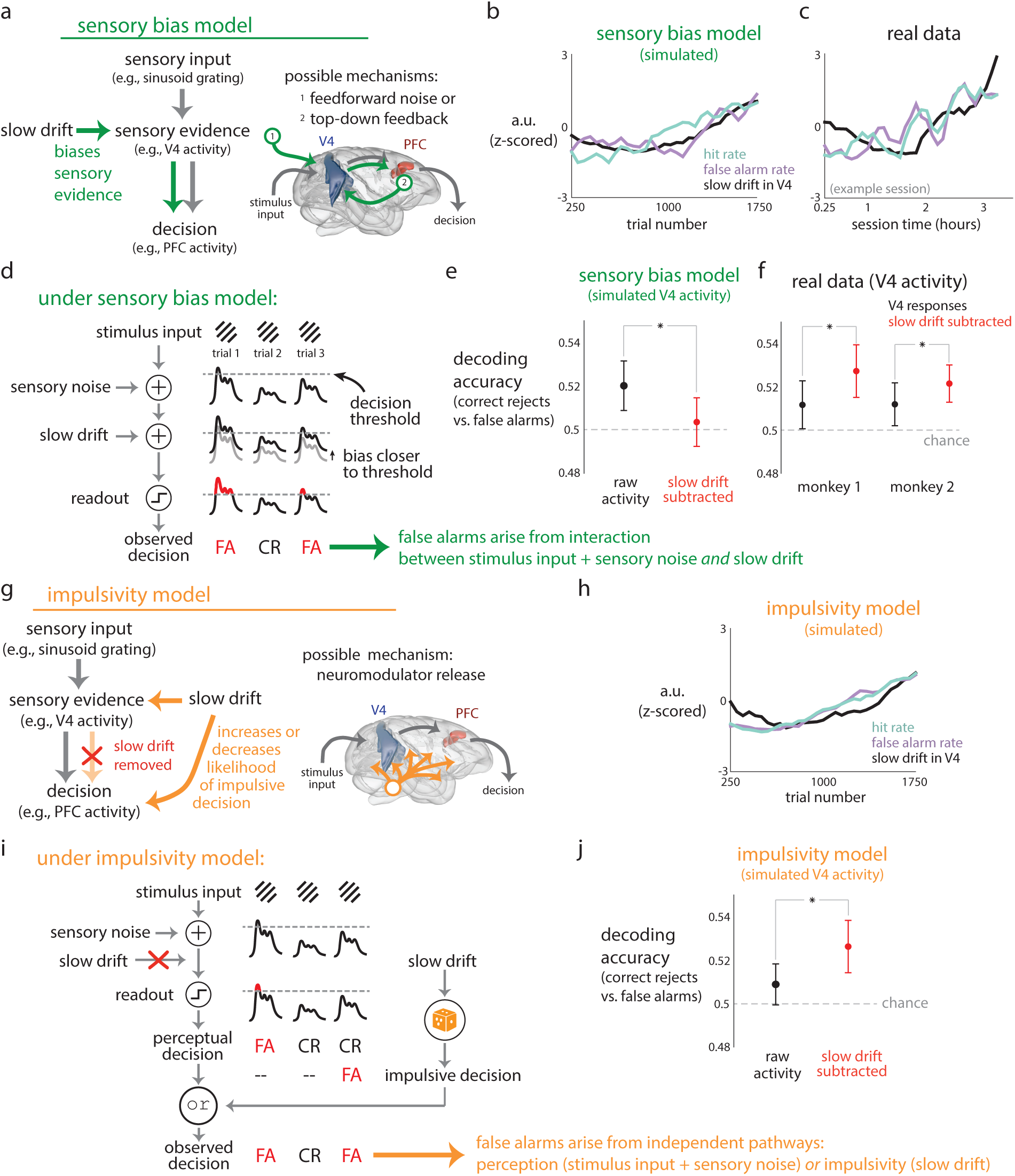
Two models of how the slow drift could influence the decision-making process. **a**. The sensory bias model. The slow drift biases sensory evidence (e.g., V4 activity), and this biased evidence is then carried to downstream areas (e.g., PFC activity). The slow drift could arise from feedforward sensory noise (e.g., slow fluctuations in LGN or V1) or from top-down feedback (e.g., a slow modulation originating from PFC) (right inset). **b**. Simulations of the sensory bias model, for which hit rate, false alarm rate, and the slow drift covary over time. **c**. The covariation of the simulated variables in **b** could also be observed in real data (an example session shown). Parameters of the model were not fit to data, and thus we do not expect an exact match of lines between panels. **d**. Under the sensory bias model, false alarms occur in two ways. First, sensory noise (e.g., feedforward noise from retina or LGN) may corrupt stimulus input (e.g., sinusoidal gratings) and push sensory evidence (e.g., V4 activity, black lines) to pass a decision threshold (dashed line), leading to a perceived change in stimulus and a false alarm (leftmost red FA). Second, the slow drift may bias sensory evidence to pass the decision threshold, also leading to a false alarm (rightmost red FA). **e**. For simulations of the sensory bias model, we measured the extent to which simulated V4 activity (with or without the slow drift) predicted false alarms (see Methods). Decoding accuracy *decreased* when the slow drift was subtracted from the simulated V4 activity (red dot below black dot, *p <* 0.002, paired permutation test). Dots indicate means, error bars indicate ±1 s.d. over 50 runs. **f**. For the real data, subtracting the slow drift from V4 activity *increased* decoding performance (red dots above black dots, asterisks correspond to *p <* 0.05, paired permutation test). Dots indicate means over sessions, error bars indicate ±1 s.e.m. **g**. The impulsivity model. The slow drift directly influences downstream areas to increase or decrease the likelihood of an impulsive decision (i.e., a saccade made independently from sensory evidence). The slow drift is present in sensory evidence (e.g., V4 activity), but downstream areas remove the slow drift from their readouts (red X). The slow drift could arise from a brain-wide release of neuromodulators (right inset). **h**. The impulsivity model reproduced the finding in real data that hit rate, false alarm rate, and the slow drift covary over time (cf. panel **c**). **i**. Under the impulsivity model, false alarms occur in two ways. First, like in the sensory bias model, sensory noise may lead to a false alarm (leftmost red FA). Second, the slow drift may increase the likelihood of an impulsive decision (orange die), leading to a false alarm (rightmost red FA). **j**. Under the impulsivity model, decoding accuracy of predicting false alarms from V4 activity *increased* when we subtracted the slow drift from simulated V4 activity (red dot above black dot, *p <* 0.002, paired permutation test), consistent with the real data (cf. panel **f**). Dots indicate means, error bars indicate ±1 s.d. over 50 runs.

To test this sensory bias hypothesis, we simulated a simple decision-making model in which V4 activity was thresholded to determine the final decision (Supplementary Fig. 9, see Methods). We biased the simulated V4 activity to be closer to or farther from a decision threshold by adding a slow drift that varied across simulated trials. We confirmed that the outputs of the sensory bias model (Fig. 6**b**) were consistent with our real data (Fig. 6**c**, representative session) in that hit rate, false alarm rate, and slow drift covaried together. Next, we developed an analysis to probe how errors occur on a trial-by-trial basis. Under the sensory bias model, false alarms occur because both sensory noise and the slow drift can push sensory evidence (i.e., V4 activity) past a decision threshold (Fig. 6**d**). Thus, subtracting the slow drift from V4 activity removes an important signal on which the final decision depends. Indeed, in our simulations, we found that subtracting the slow drift from V4 activity led to a *decrease* in how well V4 activity predicted the occurrence of false alarms within a trial (Fig. 6**e**, red dot below black dot).

To see if this effect held for real data, we performed the following analysis. We first computed how well V4 activity predicted the occurrence of a false alarm versus a correctly-rejected flash within a trial. We considered responses during a 175 ms time window between stimulus onset and saccade onset (see Methods). We found that these responses predicted false alarms above chance (Fig. 6**f**, black dots above dashed line, *p <* 0.05 for both monkeys, one sample *t*-test). Next, we subtracted the estimate of the slow drift from V4 activity, re-trained the decoder, and computed the decoding accuracy. We found that subtracting the slow drift significantly *increased* decoding accuracy (Fig. 6**f**, red dots above black dots). This result did not support the sensory bias model (compare Fig. 6**e** to **f**).

That the slow drift is a neural fluctuation that does not bias sensory evidence is not completely surprising. Previous work has reported neural fluctuations in sensory cortical neurons that are seemingly unrelated to sensory encoding but rather related to factors such as locomotion [27], arousal [64, 100], fidgeting [69], and thirst [2]. However, it is unclear how a downstream area could reliably decipher sensory evidence in the presence of constantly drifting neural activity. One intriguing possibility is that the downstream areas that read out activity from sensory areas may somehow account for these fluctuations.

Inspired by this possibility, we hypothesized that the slow drift reflects a separate decision process, independent of sensory evidence, that leads the animal to be more or less likely to make a saccade (i.e., an increase or decrease in both hit rate and false alarm rate). In previous work, similar behavioral processes have been described as urgency in a drift diffusion model [15, 34], exploration in a multisensory discrimination task [76], and impulsivity in a response inhibition framework [6]. For our purposes, we term this process “impulsivity,” which reflects the animal’s tendency to make a decision without incorporating sensory evidence (Fig. 6**g**). Under this model, the slow drift acts as an impulsivity signal, increasing or decreasing the likelihood of making a saccade independent of sensory evidence. It does this by directly influencing downstream areas that form the decision, overriding the sensory evidence. The slow drift might be attributed to a brainwide release of neuromodulators (Fig. 6**g**, right inset), consistent with our findings that V4 and PFC slowly drift together (Fig. 5). This neuromodulator could arise from brainstem nuclei and influence both sensory processing areas as well as the downstream areas in which the final decision is formed. To prevent the slow drift from interfering with the sensory readout process (i.e., what occurs in the sensory bias model), the impulsivity model removes the slow drift from its perceptual readout (Fig. 6**g**, red X). The brain may perform this removal by having a downstream area access a copy of the slow drift via the same neuromodulator that modulates V4 and subtract this modulation (e.g., through inhibition) from its sensory readout.

We simulated the impulsivity model (Supplementary Fig. 9, see Methods), and found that the slow drift produced by the model covaried with hit rate and false alarm rate (Fig. 6**h**), consistent with the real data (Fig. 6**c**). Under this model, some false alarms occur due to sensory noise (Fig. 6**i**, perceptual decision, left FA) while others occur due to impulsivity (Fig. 6**i**, impulsive decision, right FA). We next asked if subtracting the slow drift from V4 activity increased decoding performance, as observed in the real data (Fig. 6**f**). In this model, the slow drift obscures the sensory evidence in V4 activity (i.e., stimulus input + sensory noise), upon which the decision to saccade or not is based (i.e., whether the sensory evidence crosses a threshold). This is different from the sensory bias model, for which the sensory evidence comprises stimulus input + sensory noise + the slow drift. It then follows that for the impulsivity model, subtracting the slow drift from V4 activity would improve our predictions of false alarms that arise from perceptual decisions because we remove a “nuisance” variable that otherwise obscured the sensory evidence. Indeed, we found this to be the case (Fig. 6**j**, red dot higher than black dot). Thus, the impulsivity model is more consistent with the real data than the sensory bias model (cf. Fig. 6**j** and **f**).

There are two important components of the impulsivity model and, without both, the model would fail to be consistent with real data. First, without the slow drift acting as an impulsivity signal, the model would fail to have fluctuations in its behavioral output (i.e., teal and purple lines would be flat in Fig. 6**h**). Second, without the removal of the slow drift from readouts of V4 activity, the model would fail to show an increase in decoding accuracy when the slow drift is subtracted from V4 activity (i.e., red dot would not be above black dot in Fig. 6**j**). Thus, both of these components allow the impulsivity model to capture aspects of the real data that the sensory bias model cannot. In the sensory bias model, by contrast, because the slow drift is a component of the sensory evidence that influences the decision-making process, subtraction of the slow drift reduces the ability to predict behavior.

### Why is it helpful to remove slow drift from sensory activity?

Our results are more consistent with the impulsivity model in which the slow drift is removed from the perceptual decision making process. These results are based on a task in which the animals detect changes in a small range of visual stimuli (i.e., sinusoidal gratings). Thus, we next asked whether this removal process would be needed in a more naturalistic perceptual setting, which we address in the next section.

We have found evidence that a downstream area likely removes the slow drift from its readout of V4 activity. Why would the brain employ such a mechanism? One reason is that if not removed, the slow drift could corrupt sensory information encoded by V4 activity and negatively impact perception and non-impulsive decisions. Consider the following illustrative example. The responses of two V4 neurons to different natural images encode information about these images along a ‘stimulus-encoding’ axis (Fig. 7**a**, left panel, dashed black line). If the slow drift lies along an axis that is not aligned with the stimulus-encoding axis (Fig. 7**a**, middle panel, dashed black and purple lines are not aligned), the response encoding remains reliable along the stimulus-encoding axis. However, if the slow drift lies along the stimulus-encoding axis (Fig. 7**a**, right panel, dashed black and purple lines are aligned), the response encoding would be corrupted because the slow drift perturbs the responses over time independent of which stimulus is shown. Although we considered only one stimulus-encoding axis for this example, a population of neurons likely uses multiple stimulus-encoding axes, each encoding different properties of the natural images.

**Figure 7:**
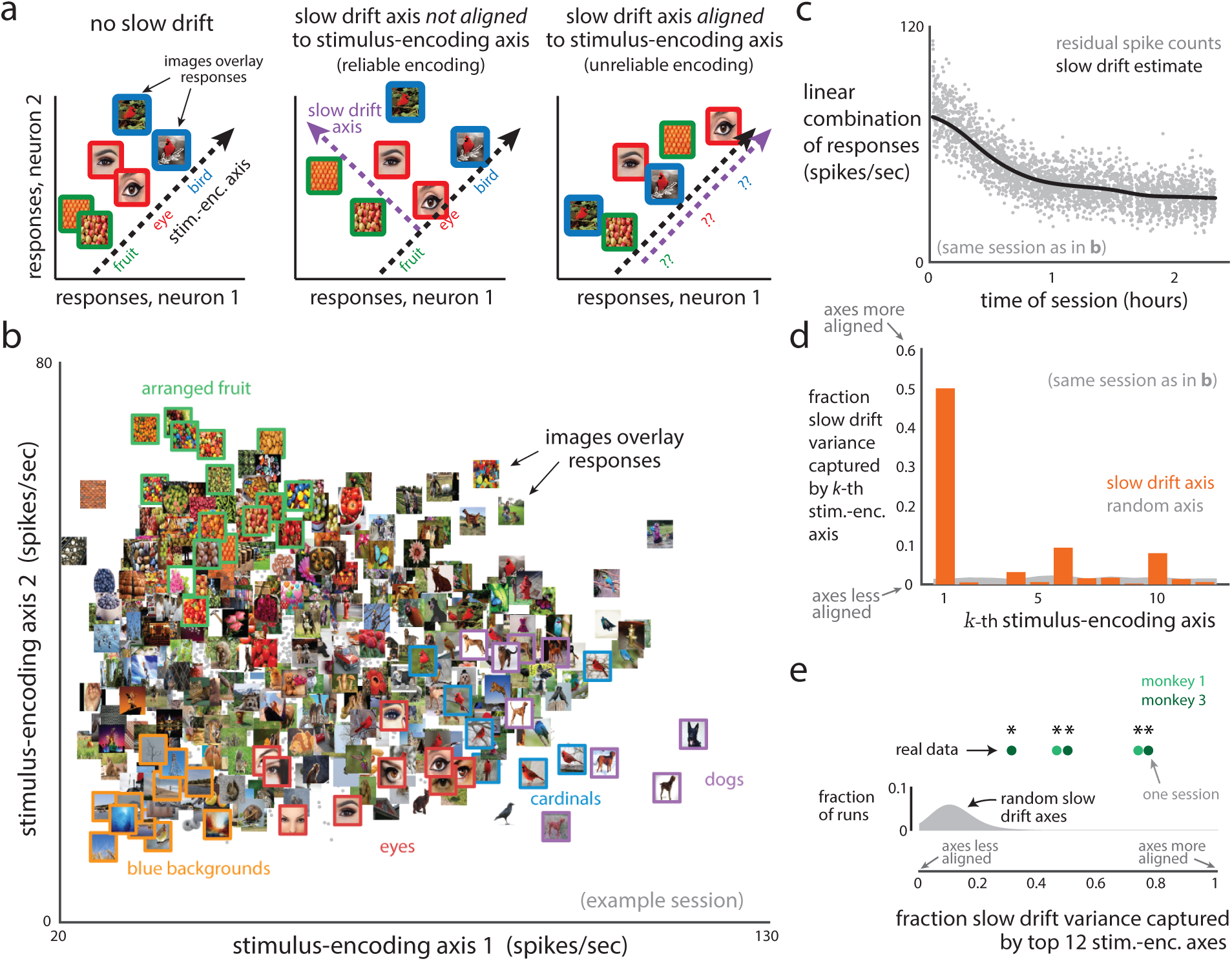
The slow drift axis is aligned to stimulus-encoding axes. **a**. Illustration of how the slow drift may influence the stimulus encoding of two V4 neurons. Without slow drift (left panel), a downstream area may read out the responses along a stimulus-encoding axis (dashed black line) and faithfully recover stimulus information (e.g., fruit, eye, bird). If the slow drift axis (middle panel, dashed purple line) is not aligned to the stimulus-encoding axis, the encoding remains reliable. If the slow drift axis is aligned to the stimulus-encoding axis (right panel), the encoding is corrupted by the slow drift and is unreliable because the slow drift displaces responses over time, independent of which stimulus is shown. **b**. Repeat-averaged V4 responses to 2,000 natural images (gray dots) along the first two stimulus-encoding axes (identified by applying PCA to the responses of 129 neurons for this session). Selected images were overlaid on top of their corresponding responses, and colored outlines denote similar images that resulted in similar responses. **c**. Linear combination of the activity of the 129-simultaneously-recorded neurons from the same session as in **b**. Same conventions as in Fig. 2**c**. Each gray dot corresponds to the residual spike counts (raw spike counts minus repeat-averaged responses) binned over each trial and projected onto the slow drift axis. These projections are then Gaussian smoothed (black line) to identify the slow drift. **d**. The fraction of slow drift variance captured by each stimulus-encoding axis (orange) for the same session as in panels **b** and **c**. A fraction closer to 1 indicates that the stimulus-encoding and slow drift axis are more aligned. The top two stimulus-encoding axes (*k* = 1, 2) correspond to the two axes in panel **b**. The top 12 stimulus-encoding axes captured 77% of the slow drift variance, significantly higher than if the slow drift lay along a randomly-chosen axis (gray, captured on average 9% of the slow drift variance, *p <* 0.002, proportion of 500 runs whose fraction was greater than that of the slow drift). The small fraction observed for the randomly-chosen axes stems from the fact that a random axis is unlikely to align with 12 other axes in a high-dimensional space (where the dimensionality is equal to the number of neurons, 129 in this case). **e**. Summary of results (five sessions from two monkeys, 75 to 129 neurons for each session). The fraction of slow drift variance captured by the top 12 stimulus-encoding axes (green dots) was significantly greater for each session than if the slow drift lay along a random axis (*p <* 0.002, proportion of 500 runs whose fraction was greater than that of the slow drift, gray distribution corresponds to 500 runs for session with smallest fraction, distributions for other sessions were similar).

Given recent theoretical studies that predict most neural fluctuations do not corrupt stimulus encoding [67, 43, 49], we predicted that the slow drift would not lie along any of the stimulus-encoding axes. We tested this hypothesis in a separate set of experiments in which monkeys performed a fixation task while viewing many natural images (see Methods). We applied PCA to the trial-averaged responses to identify axes along which they varied the most [21, 100], and defined these axes as the *stimulus-encoding axes*. The responses along the top two stimulus-encoding axes appeared to encode complex features of the images, as nearby responses corresponded to similar high-level features (Fig. 7**b**, images of blue backgrounds, eyes, cardinals, dogs, arranged fruits, etc.).

Next, we asked whether V4 activity during this fixation task contained a slow drift as in the change detection task (cf. Fig. 2). Using the same approach to identify the slow drift as before, we found a slow drift in the V4 responses (Fig. 7**c**). This suggests that the presence of the slow drift was not dependent on the type of task (change detection vs. active fixation) nor stimulus set (sinusoidal gratings vs. natural images), but rather occurs across multiple contexts.

We then tested if the slow drift axis was aligned with the top stimulus-encoding axes. We computed the fraction of slow drift variance (i.e., variance of the slow drift across time within a session) captured by each of the top stimulus encoding axes (Fig. 7**d**, orange), and compared that to a chance level for which the slow drift lies along a random axis (Fig. 7**d**, gray). We found that the total fraction of slow drift variance captured by the top 12 stimulus-encoding axes was significantly greater than the fraction expected if the slow drift lay along a random axis (Fig. 7**e**, see Supplementary Fig. 10 for individual sessions). These results indicate that a slow drift is present in V4 activity across multiple experimental contexts and lies along axes that contain stimulus information likely used by down-stream areas. If not removed, the slow drift could corrupt this stimulus information (Fig. 7**a**, right panel, unreliable encoding). This supports the notion that downstream areas account for the slow drift and remove it from their readouts of V4 activity in order to preserve reliable encoding. Thus, these results are consistent with the impulsivity model, of which an important component is this removal of the slow drift. Indeed, removing some of the neural fluctuations in readouts of visual cortical activity is likely one of the mechanisms by which the brain maintains stable and robust representation of the visual environment.

## Discussion

While monkeys perform perceptual decisions, there are slow changes in their behavior over the course of tens of minutes unrelated to task conditions, reflecting a fluctuating internal state. We discovered a neural signature of this internal state in the form of a slow drift in neural activity that covaried over time with the slow fluctuations of behavior. We found that the slow drift is a brainwide signal, present in both V4 and PFC. We further asked what role the slow drift plays in the decision-making process, and found that the model most consistent with our data was one in which the slow drift is removed from readouts of sensory evidence but still influences the ultimate decision as an impulsivity signal. Our work thus reveals that much of the apparent “noise” in the responses of cortical neurons is in fact a neural signature of a fluctuating cognitive factor, evident only when considering the temporal sequence of trials within each session.

Many studies have reported large, ongoing fluctuations in neural activity that are independent of sensory stimulation [3, 37, 64, 49]. These fluctuations evolve on different timescales, from hundreds of milliseconds [3, 73, 103, 77, 8, 104] to seconds [80, 52, 100] to minutes or longer [4, 63, 74]. Studies have found that these fluctuations correlate with behavior, including pupil diameter [70, 103, 63, 42, 24, 59, 74, 100], locomotion [27, 81], attention [77, 96, 66, 38], task difficulty [86] and learning [13, 71]. In addition, these fluctuations may have a computational purpose, such as changing the structure of noise correlations to improve the fidelity of stimulus encoding [18, 65, 8, 33], passing information from one brain area to another [85, 87, 88, 95, 89, 93], or priming neurons to certain features of the stimulus [72, 9].

However, it is unclear if every fluctuation is useful to downstream processing, and it stands to reason that the brain might have developed mechanisms to deal with such internal “noise”. One possible mechanism for noise removal is the pooling of neurons that encode redundant information in order to reduce trial-to-trial variability by taking an average over neurons [111]. A second possible mechanism is that a downstream area may read out upstream neurons such that any trial-to-trial variability shared between neurons is cancelled out and thus ignored [91, 5, 32, 67, 49]. Such a mechanism might restrict the “noise” to be along dimensions in population activity space not read out by downstream regions, as has been proposed in the motor system [44, 26, 98] and oculomotor system [46], as well as in economic choice evaluation [108]. Our observation that the slow drift aligns well with stimulus-encoding dimensions suggests this mechanism could not elude the “noisy” fluctuations of the slow drift. Our work suggests yet a third type of noise removal, whereby downstream areas remove noisy neural fluctuations from their readouts. This removal could be carried out by a downstream area accessing an independent copy of the slow drift and subtracting out this copy via either inhibition [107] or dividing the readout by this copy via normalization [12]. This copy may be accessed through the presence of a neuromodulator or by a downstream area keeping a running estimate of the slow drift in working memory [16]. Our work points to potential mechanisms to remove “noise” as an important area of focus in future studies on the impact of variability on perception and behavior.

The observed correlation between hit rate and false alarm rate (Fig. 1**d**) implies that shifts in criterion occurred (i.e., shifts of the decision threshold in signal detection theory). Previous work has shown that activity in PFC [55] but *not* V4 [54] is associated with shifts in criterion. Here, we found that activity in both V4 *and* PFC was related to criterion (i.e., the slow drift covaried with both hit rate and false alarm rate). Although on the surface these results might appear to be contradictory, they are not. Instead, this difference can be attributed to the fact that Luo and Maunsell specifically measured a spatially-selective criterion signal, defined by their task structure, whereas we measured a global criterion shift independent of task structure. In particular, our findings suggest that a shift in criterion is not necessarily due to a bias in sensory evidence (i.e., a change in decision threshold), but instead can be explained by separate decision processes unrelated to sensory evidence, such as impulsivity. That V4 may have multiple signals that co-exist and influence behavior differently is not unexpected. Indeed, in addition to local criterion and impulsivity, neural fluctuations in V4 relate to reward expectation [7], attentional effort [30], and task difficulty [86], suggesting that V4 is part of a flexible, multifaceted circuit.

What is the source of the slow drift? Given that the slow drift is a brain-wide signal that covaries with pupil diameter (Fig. 3**b**), it is conceivable that the slow drift arises from the release of neuromodulators throughout the brain. One candidate neuromodulator is norepinephrine, which is distributed by the locus coeruleus (LC) to many different brain areas on a similar timescale as that of the slow drift [4, 64, 42]. It has also been proposed that the LC modulates arousal, as the activity of LC neurons has been linked to behavioral variables that reflect arousal, such as pupil diameter [63, 42, 53] and task performance [4, 25]. Another candidate neuromodulator is acetylcholine, which is released by the basal forebrain and has been shown to relate to pupil diameter and locomotion [62, 28, 64, 110]. Further experiments that include electrophysiological recordings from relevant nuclei and experimental manipulation of the levels of different neuromodulators are needed to identify the source of the slow drift.

Our findings raise an interesting conundrum: Why does the brain modulate visual cortical activity (i.e., the slow drift) and then remove this signal downstream, rather than have no modulation of visual cortical activity at all? Some studies have suggested that neural fluctuations in visual cortex enhance the fidelity of stimulus encoding by increasing firing rates and reducing noise correlations [85, 64, 8, 33, 71], or by integrating sensory evidence with feedback from higher cortical areas [72, 9, 100]. However, the fluctuations observed in these studies typically act on faster timescales (tens to hundreds of ms) than the slow drift (tens of minutes), and may be a result of mechanisms similar to those of spatial attention [37, 8, 71]. Instead, the slow drift operates on a timescale detrimental for processing stimuli that change rapidly. One possible explanation for the existence of the slow drift and its internal removal is that through evolution, the underlying neural mechanisms governing perception evolved in a coordinated manner with the mechanisms that govern the animal’s internal state in order to achieve high fidelity of stimulus readout while allowing for brain-wide releases of neuromodulators [84]. The slow drift may also be closely connected to the global fluctuations observed in sleep [99, 64]. For example, the slow drift may represent a fluctuating state between high and low levels of alertness, with corresponding differences in neuromodulation.

As studies continue to investigate richer, more naturalistic behavior and the underlying cognitive influences, there has been an increasing use of terms such as arousal, motivation, effort, urgency, impulsivity, etc. It is often unclear which label to place on a particular observed change in behavior and/or neural activity. Regardless of the label applied to our data, the slow drift we identified represents a substantial change in neural activity associated with profound fluctuations in behavior during perceptual decisions. This signal was most evident when considering how neurons change their activity together, accounted for a large fraction of the apparent noise in neural activity from trial to trial, and could be found only when we considered the time ordering of trials within a session. Our observation that slow drift is a widespread signal in cortex means that recognizing and accounting for it may be critical in any study that attempts to link cortical neural activity to behavior.

## Materials and Methods

### Electrophysiology

Experimental details have been described previously [97]. Briefly, we recorded extracellular activity from populations of V4 and prefrontal cortex (PFC) in two male, awake, head-fixed rhesus monkeys (*Macaca mulatta*). After each animal was trained to perform an orientation-change detection task, we chronically implanted two 96-electrode arrays (Blackrock Microsystems; 1 mm in electrode length, 400-*µ*m spacing in a 10 *×* 10 grid). For monkey 1 (‘Pe’), we implanted in right V4 and right PFC (on the prearcuate gyrus, dorsolateral area 8Ar), and for monkey 2 (‘Wa’), we implanted in left V4 and left PFC. Experimental procedures were approved by the Institutional Animal Care and Use Committee of the University of Pittsburgh and were performed in accordance with the United States National Research Council’s Guide for the Care and Use of Laboratory Animals.

Voltage signals were spike sorted with semi-supervised sorting algorithms [94] and visually inspected using custom MATLAB software [45], taking into account waveform shapes and inter-spike interval distributions (https://github.com/smithlabvision/spikesort). Our data consisted of both well-isolated single units and multi-units, and we refer to each unit as a “neuron”. After applying rigorous spike waveform controls (see below), each session had 7-54 recorded V4 neurons (24-54 for monkey 1 and 7-35 for monkey 2) with a median of 32 V4 neurons (40, 20 neurons for monkeys 1, 2). For recordings in PFC (for which the same spike waveform controls were applied), each session had 41-93 recorded PFC neurons (41-93 for monkey 1 and 41-85 for monkey 2) with a median of 60 PFC neurons (60, 58 neurons for monkeys 1, 2).

### Orientation-change detection task

We trained each animal to perform an orientation-change detection task in which each trial comprised a sequence of flashes, where each flash had two Gabor patch stimuli (one presented in each visual hemifield). After fixating, the animal was rewarded with water or juice for correctly detecting a change in stimulus by making a saccade to the changed stimulus (i.e., a “hit”). Any incorrect decisions, such as breaking fixation, missing a changed stimulus, or making a saccade to an unchanged stimulus, resulted in no reward and a 1 second time out before the next trial began. The display was a gamma-corrected flat-screen cathode ray tube monitor positioned 36 cm from the animal’s eyes with a resolution of 1024*×*768 pixels, refreshed at a frame rate of 100 Hz. The background of the display was 50% luminance (gray). The gaze of the animal was tracked using an infrared eye tracking system (EyeLink 1000; SR Research, Ottawa, Ontario), and monitored online by experimental control software to ensure fixation within ∼1° of the central fixation point throughout each trial. Any trials in which the animal broke this fixation window without regard to the task (e.g., saccades that did not end in one of the two stimulus locations) were removed from our analyses.

#### Stimulus details

Presented visual stimuli were achromatic, drifting Gabor patches scaled and positioned to roughly cover the aggregate receptive fields (RFs) of the recorded V4 neurons. For monkey 1, Gabor stimuli had a diameter of visual angle 7.02°, and were centered 7.02° below and 7.02° to the left of fixation. For monkey 2, stimuli had a diameter of 4.70° and were centered 2.35° below and 4.70° to the right of fixation. Gabor stimuli were full-contrast with a spatial and temporal frequency that elicited robust responses from the population overall (i.e., not optimized for any particular neuron). For monkey 1, frequencies were 0.85 cycles/° and 8 cycles/s. For monkey 2, frequencies were 0.85 cycles/° and 7 cycles/s. During the task, we presented a Gabor stimulus at the estimated RF location and simultaneously at the mirror-symmetric location in the opposite hemifield.

#### Trial details

The trial structure of the task was as follows. At the start of the trial, the animal fixated at a centrally-located yellow dot (0.6° in diameter) on a blank, isoluminant screen. Each trial comprised a sequence of flashes, where each flash was a 400 ms presentation of two Gabor stimuli, one in each visual hemifield, followed by a blank screen lasting for 300-500 ms (uniformly distributed). For each trial, the orientation angle of one stimulus was randomly chosen to be 45° or 135°, and the orientation of the stimulus in the opposite hemifield was orthogonal (either 135° or 45°, respectively). Subsequent flashes in the sequence each had a fixed probability (30%, 40% for monkeys 1, 2) of the change in orientation of one of the stimuli (i.e., the “target”). Stimulus sequences continued until either the animal made an eye movement (i.e., a “hit” or a “false alarm”) or the animal remained fixating for 700 ms after a target appeared (i.e., a “miss”). The average sequence length (i.e., the number of flashes per trial, determined either by when a target was presented or when the animal made a false alarm) was 2.9, 2.7 flashes for monkeys 1, 2. The longest sequence length was 20, 14 flashes for monkeys 1, 2. For most of our analyses, we consider the stimulus flashes that occur after the first stimulus flash and before the final stimulus flash in a trial’s sequence (i.e., sequence positions 2, …, *M* − 1 for a sequence with *M* flashes), which we designate as *sample stimuli*.

#### Cueing blocks of trials to probe attention

To probe the effects of spatial attention, trials were blocked in an alternating fashion. Within a “cue-in” block, 90% of the stimulus changes (“valid” trials, randomly chosen) occurred for the stimulus inside the RFs of the recorded V4 neurons, while the remaining 10% of stimulus changes (“invalid” trials) occurred in the opposite visual hemifield. For valid trials, the orientation change was randomly chosen to be 1°, 3°, 6°, or 15° in either the clockwise or anti-clockwise direction. For invalid trials, we restricted the orientation change to be only 3° randomly in either the clockwise or anti-clockwise direction. This restriction was necessary in order to provide enough trials to reasonably estimate the animal’s rate of detecting stimulus changes for these infrequent trials. The other type of block was “cue-out,” which had the same task structure and percentages as cue-in blocks except that for valid trials of cue-out blocks, the stimulus change occurred outside the RFs of the recorded V4 neurons.

Each block lasted until the animal made 80 correct detections in that block, at which point the type of block (cue-in or cue-out) switched. To alert the animal to the direction to attend in each block, each block began with a set of initial trials in which only the valid stimulus was presented (with no stimulus in the opposite hemifield). These initial trials lasted until the animal correctly detected 5 orientation changes, after which pairs of stimuli were presented for the remainder of the block. These initial trials were excluded from all analyses. Block type (left or right cue) alternated within a session, with the first block type counterbalanced across sessions. The average number of trials within each block was 179, 224 trials for monkeys 1, 2. Monkey 1 completed 25 sessions of the experiment; monkey 2 completed 24 sessions. After excluding sessions for equipment failure (for 2 sessions, the photodiode signal used to align eye-tracking and neural data was unexpectedly lost) and sessions with 5 neurons or less (4 sessions), we analyzed 24 sessions completed by monkey 1, and 19 sessions completed by monkey 2.

We reported how the animal’s sensitivity *d*^’^ increased for larger changes in grating orientation (Fig. 1**b** and Supplemental Fig. 1). From the hit rate and false alarm rate estimates (see below), we also computed sensitivity as *d*^’^ = *ϕ*(hit rate) *− ϕ*(false alarm rate), where *ϕ*(*·*) is the inverse cumulative distribution function of the Gaussian distribution, following previous studies [56, 54]. We note that our task is not a true two-alternative forced choice task, as for each stimulus flash the animal could choose to make a saccade to one of two targets or remain fixating. However, because the animal rarely made a false alarm to the uncued location or chose the unchanged stimulus when the target was presented (less than 1% of trials), we assume that the choice of the animal follows statistics similar to those of a two-alternative forced choice task, and thus we report sensitivity.

### Quantifying the slow fluctuations in behavior

We analyzed two common metrics of the animal’s behavior: hit rate and false alarm rate. *Hit rate* was defined as the number of correct saccades toward a target divided by the total number of times a target appeared. *False alarm rate* was defined as the number of saccades toward a sample stimulus (i.e., not a target) divided by the total number of presented sample stimuli (excluding the initial stimulus presentation, which was never a target). We took running rate estimates in 30 minute windows, shifting the window in 6 minute increments. The long duration of the time windows (30 minutes) was necessary to ensure reliable estimates of rate over a relatively small number of trials (∼300 trials per 30 minutes; some periods had much fewer trials due to the animal briefly resting). We computed correlations between hit rate and false alarm rate over time within a session (Fig. 1**d**), and compared them to correlations when we shuffled rates across sessions (within-session time points remained unshuffled). We truncated longer sessions to have the same number of time points as the shorter sessions (discarding any time points that occur after the shorter session ended).

To assess the magnitudes of the slow fluctuations in hit rate and false alarm rate, we compared their absolute changes relative to the shifts in hit rate and false alarm rate due to spatial attention. To measure the behavioral effect size of attention, we computed the hit rate separately for valid and invalid trials across all blocks for each session, and took the difference between the two. To measure the behavioral effect size of the slow fluctuations, we computed the hit rate for each block, and took the maximum difference across all pairs of blocks for that session. We measured differences in false alarm rate in the same manner. For this comparison, we only considered trials for which a stimulus change of ±3°occurred or would occur if the animal had not false alarmed. This was because invalid trials had stimuli that only changed ±3°.

Next, we assessed the timescale of these slow fluctuations in behavior. We re-estimated hit rate and false alarm rate as above except shifting windows in 1 minute increments (instead of 6 minute increments) to allow for more time points. Then, we performed Gaussian smoothing on the running estimates with standard deviations (i.e., timescales) ranging from 1 minute to 60 minutes in 1 minute increments. For each candidate timescale, we computed a cross-validated *R*^2^ based on leaving out randomly-chosen time points for each fold (10 folds in total) and then predicting the value of each held-out point by taking a Gaussian weighted average of its neighbors. We found the timescale that maximizes this *R*^2^. We then increased the timescale until the *R*^2^ dropped to 75% of the peak *R*^2^. We define this as the timescale of the slow fluctuations in behavior. We provide intuition this approach in Supplemental Fig. 11. We confirmed that the estimated timescales were similar for different durations of time windows (e.g., 20 minutes instead of 30 minutes).

### Estimating slow drift in neural activity

To estimate the slow drift in V4 and PFC activity, we first computed residual spike counts by taking the spike counts of each repeat of the same stimulus (binned in a 400 ms epoch starting 50 ms after stimulus onset) and subtracting the mean spike counts averaged across repeats of that stimulus orientation (either 45° or 135°). We then took a running average of the residual spike counts in a 20 minute time window, where each window was offset in time from the previous window by 6 minutes. We chose a 20 minute time window (as opposed to the 30 minute window chosen for the running estimates of behavioral variables) because here we could reliably estimate the slow drift with a smaller window. However, different values of the time window width and window offset yielded similar results. We applied principal component analysis (PCA) to the running average of residual spike count vectors (20 to 40 vectors per session, where the length of each vector equaled the number of neurons), and defined the unit-length vector of weights (i.e., loadings) of the top principal component (PC) as the *slow drift axis*. We found that on average the slow drift axis explained ∼70% of the variance of the running average of residual activity (69%, 77% for monkeys 1, 2). We then projected the residual spike counts (400 ms bins) onto this axis. Next, we performed Gaussian smoothing (with a timescale of 9 minutes; different values yielded similar results), and defined the Gaussian-smoothed projections as the *slow drift*. The specified time window width, window offset, and smoothing timescale were chosen in a reasonable range and not optimized. We chose to use PCA instead of factor analysis (FA) [109, 106], because the length of the time windows (20 minutes) likely averaged away most Poisson-like response variability that would otherwise be better described by FA. Using FA instead of PCA yielded results similar to those presented here.

### Aligning the V4 slow drift across sessions

The sign of the slow drift was arbitrary because PCA identifies the orientation of the slow drift axis in the population activity space up to a 180° rotation. Thus, without an alignment procedure, the sign of the correlation between the V4 slow drift and a behavioral variable was arbitrary. To combine the correlations across sessions (Fig. 3**b**), we needed a way to align the slow drift axis that was consistent across sessions. One possibility was to flip the sign of the slow drift such that the sum of its axis weights was positive. However, this procedure did not account for the possibility that different sessions may have different proportions of recorded neurons with positive or negative weights (and hence making this flipping procedure sensitive to which neurons happened to be recorded in that session). Instead, we adopted a different procedure that established an absolute reference across sessions based on the sample stimuli of the experiment (i.e., grating orientation angles of 45° or 135°). The reasoning behind this alignment procedure was that the slow drift was likely aligned to the stimulus representation of the population of neurons, and because our arrays were implanted in one location and remained there, and the response properties of nearby neurons are similar and likely remain constant over time, this alignment was similar across sessions. We aligned the orientation of the slow drift axis such that the projections of the mean spike counts of the two sample stimuli along the slow drift axis always yielded a higher projection value for the 45° stimulus than that for the 135° stimulus. Importantly, this alignment was independent of any observed behavioral variables, and thus any correlation between the slow drift and a behavioral variable over the session cannot be due to this alignment procedure. This alignment procedure was used in Figure 3**b**, Supplemental Figure 7, and Supplemental Figure 8.

There were a small number of sessions in which the population responses to the two orientation angles were not well differentiated. In these cases, the alignment procedure did not produce a meaningful result. To identify and remove those sessions which would have otherwise reduced our ability to observe the relationship between neural effects and behavior, we decoded the stimulus identity (45° or 135°) from the population responses using a linear SVM in a cross-validated manner. When computing the correlations between the slow drift and behavioral variables (Fig. 3**b**), we included only sessions with a decoding accuracy greater than 55% (21/24 sessions for monkey 1, 18/19 sessions for monkey 2). Note that we could have flipped the slow drift such that the higher projection value was for the 135° stimulus instead of the 45° stimulus. This would result in a change of sign with the same magnitudes for all correlations (i.e., in Fig. 3**b**, correlations for hit rate, false alarm rate, and pupil diameter would be negative, and correlations for trial duration and reaction time would be positive).

### Controlling for neural recording instabilities

The slow drift could have arisen from non-neural sources, such as neural recording instabilities. Such instabilities may cause the spike waveforms to gradually change shape throughout a session, and thus affect our spike sorting procedure. For example, if the spike waveform changes over time, spike sorting may be more likely to miss spikes at different times of the session, leading to a slow drift. To ensure that the observed slow drift was not due to neural recording instabilities, we used the following two criteria for the inclusion of a V4 or PFC neuron in our analyses.

First, we included only neurons whose spike waveform was stable throughout the session. To do so, we divided the session into 10 non-overlapping time bins of equal size, and computed the mean spike waveform in each bin as well as the mean spike waveform across the entire session. Then, we computed the squared norm of the difference between each bin’s mean waveform and the overall mean waveform. The *percent waveform variance* was computed as the largest squared norm across bins divided by the variance of the overall mean waveform. Example mean spike waveforms and their corresponding percent waveform variances are shown for V4 neurons in Supplementary Figure 3 and for PFC neurons in Supplementary Figure 12. We removed neurons from our analyses whose percent waveform variance was above 10%. We confirmed that the remaining neurons with the largest percent waveform variances were not more likely to contribute to the slow drift than other neurons (Supp. Fig. 3c and Supp. Fig. 12).

Second, we excluded any neurons for which there was an abrupt change in activity during the session. For each neuron, we divided the session into 20 non-overlapping time bins of equal size, and computed the mean spike count within each bin. If the change in activity between any two consecutive time bins was greater than 25% of the maximum activity of that neuron (i.e., the largest of the mean spike count values across the 20 bins), the neuron was excluded from all analyses.

### Measuring behavioral variables and relating them to the slow drift

To characterize the animal’s behavior, we measured hit rate and false alarm rate (described previously), as well as the following three quantities: 1. *Pupil diameter* was the mean diameter of the pupil (arbitrary units) during each stimulus flash measured with the EyeLink eye-tracking software (EyeLink 1000; SR Research, Ottawa, Ontario). 2.

*Trial duration* was the total amount of time taken to complete a trial. Because the animal could choose to saccade to an unchanged stimulus flash (thus ending the trial), the animal’s false alarm rate influenced trial duration. We computed trial duration by taking the difference between the stimulus onset of the initial flash and saccade onset (or, when no saccade, the end time of the final flash). 3. *Reaction time* was computed only for correct trials (i.e., “hits”) as the time between the onset of the changed stimulus in the final flash and saccade onset (the time at which the eyes left the fixation window).

A running mean of each behavioral variable was taken (except pupil diameter, for which a running median was taken) with 30 minute overlapping windows in 6 minute intervals. The long length of the time window was necessary to have reasonable estimates of hit rate and false alarm rate. For fair comparison between behavioral variables and the slow drift, we also estimated the slow drift as a running mean of residual spike counts along the slow drift axis with the same time window and intervals as those for the behavioral variable estimates (i.e., Gaussian smoothing was not performed here).

We used the running estimates of the behavioral variables and slow drift for two comparisons. First, to assess whether the slow drift and behavioral variables showed similar time courses during a session, we computed the correlation over time between the slow drift and each behavioral variable (hit rate, false alarm rate, pupil diameter, trial duration, and reaction time). To compute the “shuffled” distributions of correlations between the slow drift and behavioral variables, we randomly shuffled slow drifts across sessions and re-computed correlations (200 runs), truncating the longer time series by discarding time points after the shorter times series finished. Second, we assessed if sessions for which the slow drift strongly varied (i.e., had a large magnitude) corresponded to sessions for which the behavioral variables also strongly varied. We measured the magnitude of the slow drift as its variance over time within a session. We normalized the slow drift by the number of neurons for each session to account for differences in the number of neurons across sessions. The variances of pupil diameter across sessions were not comparable because the eye-tracking software output arbitrary units of pupil diameter, and those units changed somewhat from session to session as the tracker position was adjusted. Thus, although values of pupil diameter were comparable within a session, values of pupil diameter were *not* comparable across sessions. We did not include the variance of pupil diameter in our results (Fig. 3**c** and Supp. Fig. 7).

### Measuring whether neural responses vary more strongly with the slow drift versus attention

To understand how large were the neural activity changes related to the slow drift, we compared it to the size of activity changes related to spatial attention. For spatial attention, we first defined the *attention axis* as the axis in population activity space that connects the mean response on cue-in trials to that on cue-out trials [19, 61, 97]. The mean is taken over repeats of the same sample stimulus for a given cue condition. To assess the size of attention’s effect on spike counts for a given session, we computed the mean response along the attention axis for each block of trials (Fig. 4**b**, orange and green lines), and took the variance across blocks (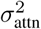). To measure the size of the slow drift, we computed the mean slow drift value for each block (Fig. 4**a**, black lines), and computed the variance of these mean values across the blocks for each session (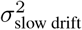). We compared the relative sizes of the slow drift and attentional effects by taking the ratio 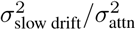. A ratio greater than 1 indicates that the size of the slow drift was larger than the modulation in responses due to attention. Ratios were computed separately for sample stimuli with 45° or 135° orientations, and results were aggregated across orientations and sessions (Fig. 4**c**). To control for the possibility that the attention axis was aligned to the slow drift axis (and hence the slow drift could leak into the estimates of the attentional effect), we subtracted the estimate of the slow drift from responses before projecting them along the attentional axis. For most sessions, we found that the slow drift axis and the attention axis were largely unaligned (i.e., this subtraction procedure often did not change the value of 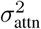). Note that 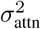 included the variance between cue-in and cue-out blocks (i.e., between orange and green lines) as well as the variance within cue-in and cue-out blocks separately (i.e., between orange lines and between green lines). This latter variance is necessary to ensure a fair comparison with the size of the slow drift but implies that 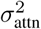 is an overestimate. Thus, the true ratios are likely to be even larger than reported in Fig. 4**c**.

### Comparing the slow drift between brain areas V4 and PFC

To compare the correlation between the V4 slow drift and PFC slow drift, we computed the correlation between the slow drift identified for each brain area separately. Then, we took the absolute value of the correlation because its sign is arbitrary. We compared the measured correlations to those computed when the PFC slow drifts were randomly shuffled either within sessions (which assumes no slow drift in PFC activity, see Fig. 5**b**, ‘shuffled’) or across sessions (which assumes a slow drift in PFC activity that is not related to the slow drift in V4 activity). For the former, correlations were expected to be larger than 0 (Fig. 5**b**, median *|ρ|* = 0.50) because we took absolute correlations. For the latter, if two shuffled sessions differed in length, we truncated the longer session by removing time points that occurred after the end of the shorter session. Again, because we took absolute correlations, correlations of the shuffled-across-sessions data were expected to be high (measured to be median *|ρ|* = 0.87) but were still significantly less than the correlations for the real data (Fig. 5**b**, median *|ρ|* = 0.96, *p <* 0.002, permutation test). We also computed the correlation between the magnitudes of V4 and PFC across sessions (Fig. 5**c**). Magnitude was defined as the variance of the slow drift (either in V4 or PFC) within a session. We normalized the slow drift by the number of neurons to account for differences in numbers of neurons across sessions.

### Models of perceptual decision-making

We propose two models of perceptual decision-making (Fig. 6 and Supplementary Fig. 9). Both models take as input the stimulus *x*_input_ *∈ {*0, 1*}* (where *x*_input_ = 0 for a sample stimulus and *x*_input_ = 1 for a changed stimulus) and feedforward perceptual noise *∊*_noise_ *∼ N* (0*, σ*^2^), and output a decision variable *d ∈ {*0, 1*}* (where *d* = 0 represents keeping fixation and *d* = 1 represents making a saccade). Similar to the experiment, each simulated trial comprised a sequence of stimulus flashes where each flash had a 40% chance of changing from *x*_input_ = 0 to *x*_input_ = 1. In a correct trial, the model outputs decision *d* = 0 when *x*_input_ = 0, and outputs *d* = 1 when *x*_input_ = 1.

Both models also have a slow drift variable 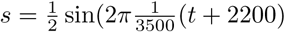, where *t* = 1, …, 2000 indicates the trial index. We chose this sine function so that *s* slowly varies over 2000 trials; other smooth functions are possible. We define *v* = *x*_input_ + *∊*_noise_ + *s* which represents V4 activity and provides sensory evidence to decision *d*. In addition to the *∊*_noise_ noise, we included another source of noise that directly influenced *d*, whereby, for each flash, there was a probability *ψ* that *d* = 1 or *d* = 0 (both equally likely), independent of any other variables in the model. This output noise reflects other internal processes in the brain from which we cannot record that may affect an animal’s decision.

#### Sensory bias model

For the sensory bias model (Fig. 6**a**), the decision is based directly on *v*, where *d* = 1 if *v >* 0.5, else *d* = 0. An increase in *s* places *v* closer to the decision threshold of 0.5, thereby increasing the chance of a false alarm. Conversely, a decrease in *s* places *v* further from the decision threshold of 0.5, thereby requiring a higher value of *∊*_noise_ to cause a false alarm.

#### Impulsivity model

For the impulsivity model (Fig. 6**g**), the slow drift *s* directly influences decision *d* independent of the sensory evidence encoded in V4 activity *v*. Mathematically, the slow drift determines an impulsivity signal *m ∼* Bernoulli(*p*(*s*)), where *p*(*s*) is the probability of saccade given the slow drift *s*. We set 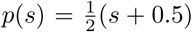, which increases linearly as *s* increases. The decision is *d* = 1 if *m* = 1. The decision is also dependent on perceptual readout *p* = *v − s*, where the slow drift has been removed. Thus, *d* = 1 if *p >* 0.5 or *m* = 1, else *d* = 0. Note that the slow drift *s* does not bias the sensory evidence because *s* is removed from the V4 activity *v*. Instead, the slow drift only influences the decision *d* by overriding the perceptual readout *p* via the impulsivity signal *m*.

#### Model simulations

We ran each model for 2,000 trials, and computed the hit and false alarm rates with running estimates over the 2,000 trials (500-trial window length, where each window was offset by 50 trials). We matched the overall false alarm rate for each model (34%) to the animals’ false alarm rates (∼30%, ∼40% for monkeys 1, 2) by choosing an appropriate value for the sensory noise parameter (*σ* = 0.35) as well as for the output noise parameter (*ψ* = 70%).

### Decoding V4 responses to predict the occurrence of false alarms

Because the outputs of both models were consistent with the finding that hit rate, false alarm rate, and the slow drift covary together (Fig. 6**b**, **c**, and **h**), we sought an analysis in which we could test a prediction that differentiated the two models. In particular, we focused on how false alarms (i.e., *d* = 1 when *x*_input_ = 0) occurred for each model. Under the sensory bias model, slow drift *s* contributes to decision *d* through the perceptual readout of V4 activity *v*. Thus, we would expect that for false alarms, *d* would be more accurately predicted from *v* than from *v − s* (i.e., subtracting the slow drift contribution from *v*). On the other hand, for the impulsivity model, slow drift *s* contributes to decision *d* through the impulsivity signal *m* but *not* the perceptual readout *p* (because *p* = *v − s*, so the contribution of *s* does not reach *d*). The slow drift *s* acts as perceptual noise in V4 activity *v*, obscuring the true perceptual signal *p* (i.e., *v* = *p* + *s*, indicating *v* comprises the perceptual signal *p* and the “noisy” slow drift *s*). In this case, we would expect that for false alarms (i.e., *p >* 0.5 when *x*_input_ = 0), *d* would be more accurately predicted from *v − s* than from *v*, which differs from the sensory bias model. We first performed a decoding analysis to verify that our expectations about these models were correct, and then performed a similar analysis for the real data to test which model’s predictions were most consistent with the real data.

#### Verifying model predictions

We performed the following decoding analysis for each model (Fig. 6**e** and **j**). First, we decoded whether or not a false alarm occurred from *v* using a threshold decoder (linear SVM). Next, we decoded from *v − s* instead of *v*. Finally, we compared the two decoding accuracies to determine if the decoding accuracy would be higher (as predicted by the sensory bias model) or lower (as predicted by the impulsivity model) when decoding from *v* than from *v − s*. Note that we decoded flashes that were either false alarms (i.e., *d* = 1 when *x*_input_ = 0) or correctly-rejected flashes (i.e., *d* = 0 when *x*_input_ = 0) that directly preceded the false alarm flashes. These correctly-rejected flashes were chosen for two reasons. First, they ensured we had an equal number of false alarm and non-false alarm flashes. Second, they forced the decoder to only consider within-trial fluctuations (i.e., *∊*_noise_) but not across-trial fluctuations (i.e., slow drift *s*). If we had instead considered across-trial fluctuations by including any correctly-rejected flash, we would not be able to differentiate between the models, as decoding accuracy for both models would be higher for *v* than for *v − s*. This is because *s* contains information about the overall probability of a false alarm during a session that could be used to predict the occurrence of a false alarm within a session (but not within a trial), and subtracting *s* would discard this information. We provide further intuition about this point in Supplementary Fig. 9.

#### Decoding analysis for the real data

We performed a similar decoding analysis for the real data as we did for each model (Fig. 6**f**). The animal’s decision was to saccade (i.e., *d* = 1) or not to saccade V4 population responses projected onto the slow drift axis (i.e., *v* is a 1-dimensional signal, see below). For *s*, we used our estimates of the slow drift of V4 activity (i.e., the population responses projected onto the slow drift axis and then smoothed). We then decoded *v* and *v − s* in the same manner as described for the models, using a linear SVM decoder with leave-one-out cross-validation.

For *v*, we processed V4 activity in the following way. Because decoding accuracy of neural activity predicting false alarms have been reported to be only slightly above chance [e.g., choice probability was found to be ∼54%, as reported in 72] and we were considering a small time window of activity (i.e., 175 ms), we leveraged the temporal information of the V4 responses. We computed spike count vectors of the V4 activity in non-overlapping 1 ms time bins (starting at stimulus onset and ending at 175 ms after stimulus onset), and performed Gaussian smoothing with a 10 ms standard deviation for each neuron. Note that compared to estimating the slow drift, the smoothing here is a different order of magnitude (10 ms versus 9 minutes) and has a different purpose. The reason for smoothing here is to account for small jitters of spike times between flashes that may occur due to the 1 ms bin widths. Trials for which a saccade occurred within 175 ms after stimulus onset were removed (less than 15% of all false alarm trials). The spike count vectors were then projected onto the slow drift axis. Thus, *v* corresponded to a 175-dimensional vector of projected spike counts (where 175 is the number of 1 ms time bins). For the slow drift *s*, we used the same procedure as described above, except that we binned activity in 175 ms time bins (instead of 400 ms time bins) and included responses to the final flashes. To obtain *v − s*, we formed a 175-dimensional vector where each element was the value of *s*.

We considered the same type of flashes (i.e., false alarms and their preceding correctly-rejected flashes) as those for the models. False alarm trials were limited to cue-in trials only, and each trial was required to have 3 or more stimulus flashes in total. To gain more statistical power, we doubled the number of data points by aggregating false alarm trials across the two orientations of stimuli (45° and 135°) in the following way. For stimulus flashes with a given grating orientation, we subtracted out the mean spike count response (the PSTH on a millisecond timescale) to the preceding correctly-rejected flash from the spike count responses to both the false alarm and correctly-rejected flashes. This ensured that the overall mean response to the correctly-rejected flash was the same for both orientations. We then analyzed all stimulus flashes together regardless of orientation.

The fact that we can weakly predict false alarms from visual cortical activity (Fig. 6**f**, ∼52%, significantly greater than the chance level of 50%, *p <* 0.05 for both monkeys, one sample *t*-test) is consistent with previous work that reported choice probabilities (∼55%) for neurons in MT, V1, V2, and V4 [72, 22, 41]. However, the observed decoding accuracies are not directly comparable to previously-reported choice probabilities because most previous work computes choice probabilities for a single neuron (here, we use a population of neurons), considers neural activity taken over large time windows (e.g., 1 second compared to 175 ms used here), and uses two-alternative forced choice tasks rather than our sequential task that required long periods of fixation. In an additional analysis, we confirmed that the resulting decoding accuracies were not a by-product of visual response adaptation due to the fact that the final flash always followed the second-to-final flash (Supp. Fig. 13).

### Determining whether the slow drift has the potential to corrupt stimulus encoding

We sought to assess whether the slow drift could corrupt stimulus encoding. To do so, we measured to what extent the slow drift lies within the “stimulus-encoding subspace”, defined as the subspace spanned by the mean responses to many natural images. The rationale is that, if the slow drift lies within the stimulus-encoding subspace, then the corrupted neural response to an image (i.e., corrupted by the slow drift) could be interpreted downstream as a different image.

To identify the stimulus-encoding subspace, we presented a large collection of natural images, which were selected using an adaptive procedure. We opted for an adaptive stimulus selection procedure because we were limited in how many images we could show per session (i.e., ∼2,000 images), and a small set of randomly-chosen images cannot fully represent the space of all possible natural images [40]. In a set of closed-loop experiments separate from those of the orientation-change detection task, we employed an adaptive stimulus selection algorithm, called *Adept*, to choose a set of 2,000 natural images (out of a candidate set of 20,000) that elicited large and diverse responses [20]. We recorded V4 activity in an adult, male rhesus macaque (*Macaca mulatta*, monkey 3, ‘Wi’) using a Utah electrode array. Surgical and electrophysiology procedures were the same as for the other two monkeys, described above. For another monkey (monkey 1), we did not use adaptive stimulus selection but rather randomly selected 550 images to show.

After choosing the collection of stimuli to show for one recording session, for subsequent sessions we presented the images in random order while the animals performed an active fixation task. The description of this active fixation task is as follows. After the animal fixated for 150 ms on a yellow dot (same display parameters as those for the orientation-change detection task) to initiate a trial, the animal remained fixating while a stimulus clip of images was presented in the RFs of the recorded V4 neurons. After the stimulus clip finished, the animal was required to maintain fixation for 700 ms, after which the fixation dot vanished and a target dot appeared ∼10 visual degrees from the location of the fixation dot. The animal received a water reward for correctly making a saccade to the target dot (within a target window of visual angle 5°). For monkey 3, each stimulus clip consisted of 10 natural images randomly selected from the 2,000 Adept-chosen natural images (no image could repeat within the same stimulus clip), and each image was shown for 100 ms. For monkey 1, each stimulus clip comprised three images (of the 550), and each image was presented for 200 ms, interleaved with 200 ms of isoluminant blank screen.

#### Identifying the stimulus-encoding axes

We identified the stimulus-encoding axes using the following procedure. We computed spike counts in 100 ms bins (for monkey 3) or 200 ms bins (for monkey 1), starting 50 ms after stimulus onset to account for the time it takes sensory information to reach V4. We defined the *stimulus-encoding axes* as the dimensions in population activity space in which the repeat-averaged responses have the greatest variance. We identified these axes using PCA, taking the weight vectors of the top *K* principal components as the top *K* stimulus-encoding axes. We did not consider responses to an image that had 5 or fewer repeats. Because repeats were not necessarily equally-spaced throughout a session, we controlled for the possibility that the slow drift “leaked” into the estimation of the stimulus-encoding axes. To do this, we first applied PCA to the repeat-averaged responses and considered all *N* PCs, where *N* is the number of neurons. For each PC, we subtracted any “slow drift” from the raw responses along that PC (where the slow drift for each PC was estimated from projected responses using Gaussian smoothing with a 9 minute standard deviation). Finally, we re-computed the repeat-averaged responses for which the slow drift was subtracted, and re-applied PCA to these responses. Results were almost identical when we did not perform this removal procedure, suggesting that little to no slow drift had leaked into the repeat-averaged responses (i.e., there were enough repeats to average out the presence of the slow drift).

#### Comparing the slow drift axis to the stimulus-encoding axes

We first estimated the slow drift of the responses to natural images. We estimated the slow drift axis using the same procedure as that for the orientation-change experiment. Note this slow drift was not necessarily the same as that removed from each stimulus-encoding axis of the repeat-averaged responses in the previous section, as this slow drift could be along an axis orthogonal to the stimulus-encoding axes. Finally, we measured the extent to which the slow drift axis was aligned to the stimulus-encoding axes by computing the fraction of slow drift variance captured by each stimulus-encoding axis. The slow drift variance was the variance of the slow drift over time within a session. A fraction close to 1 indicates that the slow drift axis largely overlaps with the stimulus-encoding axis. A fraction close to 0 indicates that the slow drift axis was close to orthogonal to the stimulus-encoding axis. For reference, we rotated the slow drift axis to a random orientation in population activity space (i.e., “random axis”), and re-computed this fraction.

### Statistical testing

Unless otherwise stated, all statistical hypothesis testing was conducted with permutation tests, which do not assume any parametric form of the underlying probability distributions of the sample. All tests were two-sided and non-paired, unless otherwise noted. We computed *p*-values with 500 runs, where *p <* 0.002 indicates the highest significance achievable given the number of runs performed. Permutation tests were performed either for differences in means or differences in medians (the latter used when outliers existed), as noted by context in the text. All correlations were performed with Pearson’s correlation *ρ*, unless otherwise stated. We computed the significance of *ρ*_actual_ of the actual data with a ‘shuffle test’ by running 500 shuffles of the samples and re-computing 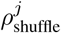 for the *j*th shuffle. The *p*-value was measured as the proportion of shuffles with 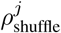 greater than or equal to *ρ*_actual_ for all *j*, for a one-sided hypothesis test. Error bars in figures represent either ±1 s.e.m. when estimating means or bootstrapped 90% confidence intervals when estimating medians, as stated. Error bars are for visualization purposes only, and not used for hypothesis testing. No statistical methods were used to predetermine sample sizes, but our sample sizes are similar to those of previous related publications [e.g., 18, 65, 54].

## Life Sciences Reporting Summary

Further information on experimental design is available in the Life Sciences Reporting Summary.

## Data availability

All relevant data are available upon request from the authors.

## Code availability

All computer code used to generate the results is available upon request from the authors. All code was written in MATLAB (The MathWorks, Natick, MA).

## Acknowledgments

The authors thank Samantha Schmitt for assistance with surgery and data collection. B.R.C. was supported by the C.V. Starr Foundation Fellowship. A.C.S. was supported by NIH K99EY025768. K.A. was supported by NSF Graduate Fellowship Grant 1747452. R.C.W. was supported by a Richard King Mellon Foundation Presidential Fellowship in the Life Sciences. B.M.Y. and M.A.S. were supported by R01MH118929, R01EB026953, and NSF NCS 1734916/1954107. B.M.Y. was also supported by NSF NCS BCS1533672, NIH R01 HD071686, NIH CRCNS R01 NS105318, and Simons Foundation 543065. M.A.S. was also supported by NIH R01EY022928 and P30EY008098.

## Author contributions

A.C.S. and M.A.S. designed the orientation-change detection task, and A.C.S. collected the data. B.R.C., K.A., and R.C.W. collected the natural image data. B.R.C., B.M.Y. and M.A.S. designed the study. B.R.C. analyzed the data and generated the figures. B.R.C. wrote the first draft of the paper, and all authors edited and provided feedback on the paper.

## Supplementary figures

**Supplementary Figure 1:**
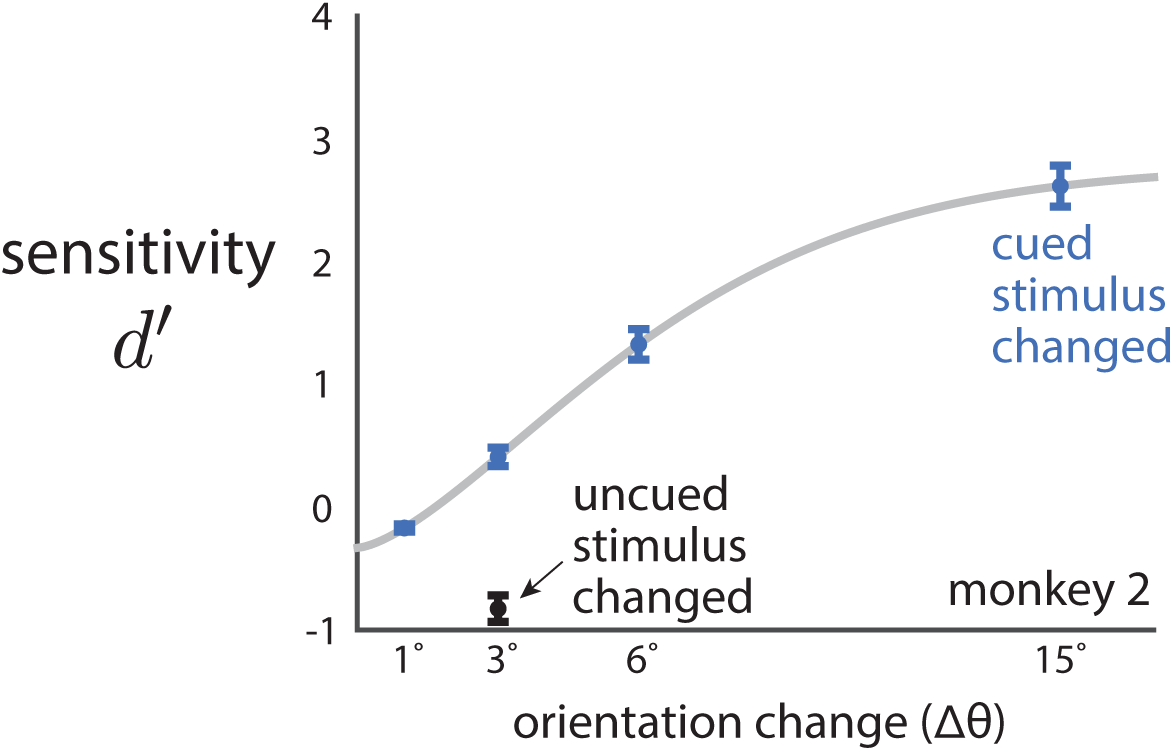
The animal better detects larger stimulus changes as well as stimulus changes that are cued than uncued. Task performance (measured as sensitivity *d*^’^) versus the change in orientation Δ*θ* (either clockwise or counterclockwise) of the cued stimulus (blue) and the uncued stimulus (black). Results shown here are for monkey 2 (see Fig. 1b for results of monkey 1). Larger Δ*θ* are easier for the animal to detect, and cued stimulus changes (Δ*θ* = 3°, blue) are easier to detect than uncued stimulus changes (Δ*θ* = 3°, black). Data are fit with a Weibull function (gray). Dots indicate means over sessions, error bars indicate ±1 s.e.m.

**Supplementary Figure 2:**
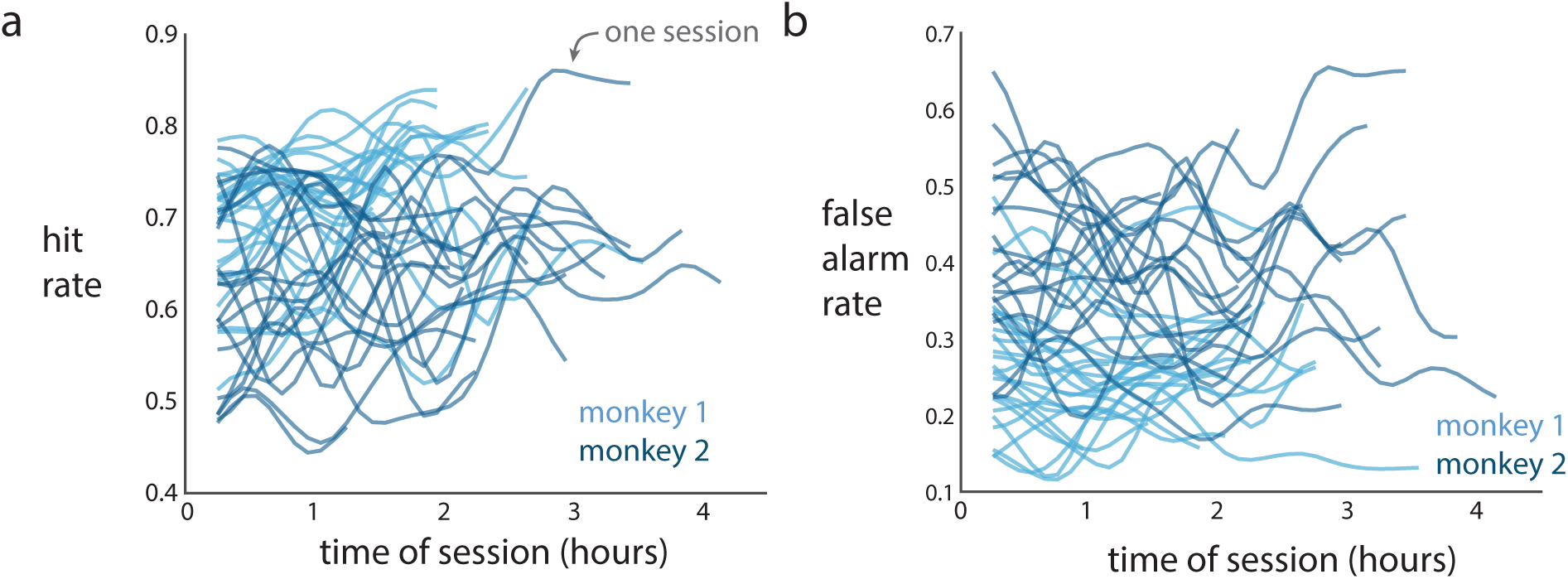
Hit rate and false alarm rate slowly change over the course of the session but not in a consistent way (e.g., all rates do not strictly increase). **a**. Hit rate, measured as the fraction of stimulus changes that were correctly detected. **b**. False alarm rate, measured as the fraction of unchanged stimulus presentations to which the animal incorrectly made a saccade. For **a** and **b**, each line is the running estimate of the rate for one session (see Methods). These results indicate that hit rate and false alarm rate did not fluctuate in a stereotypical manner (e.g., strictly increasing). This suggests that the slow changes in rates were not due to a general increase in fatigue (or satiation, etc.) over the course of the session. The reasoning is as follows. As the animal gradually becomes fatigued, the animal likely becomes less engaged in the task and more prone to “guessing”—increasing the animal’s overall likelihood to make a saccade in order to receive reward for guesses that happened to be correct. Thus, we would expect to see both hit rate and false alarm rate gradually increase throughout the session. However, hit rate and false alarm rate did not strictly increase for every session. Still, it is possible that the animal had differing periods of fatigue and high engagement throughout the session, consistent with a fluctuating hit rate and false alarm rate.

**Supplementary Figure 3:**
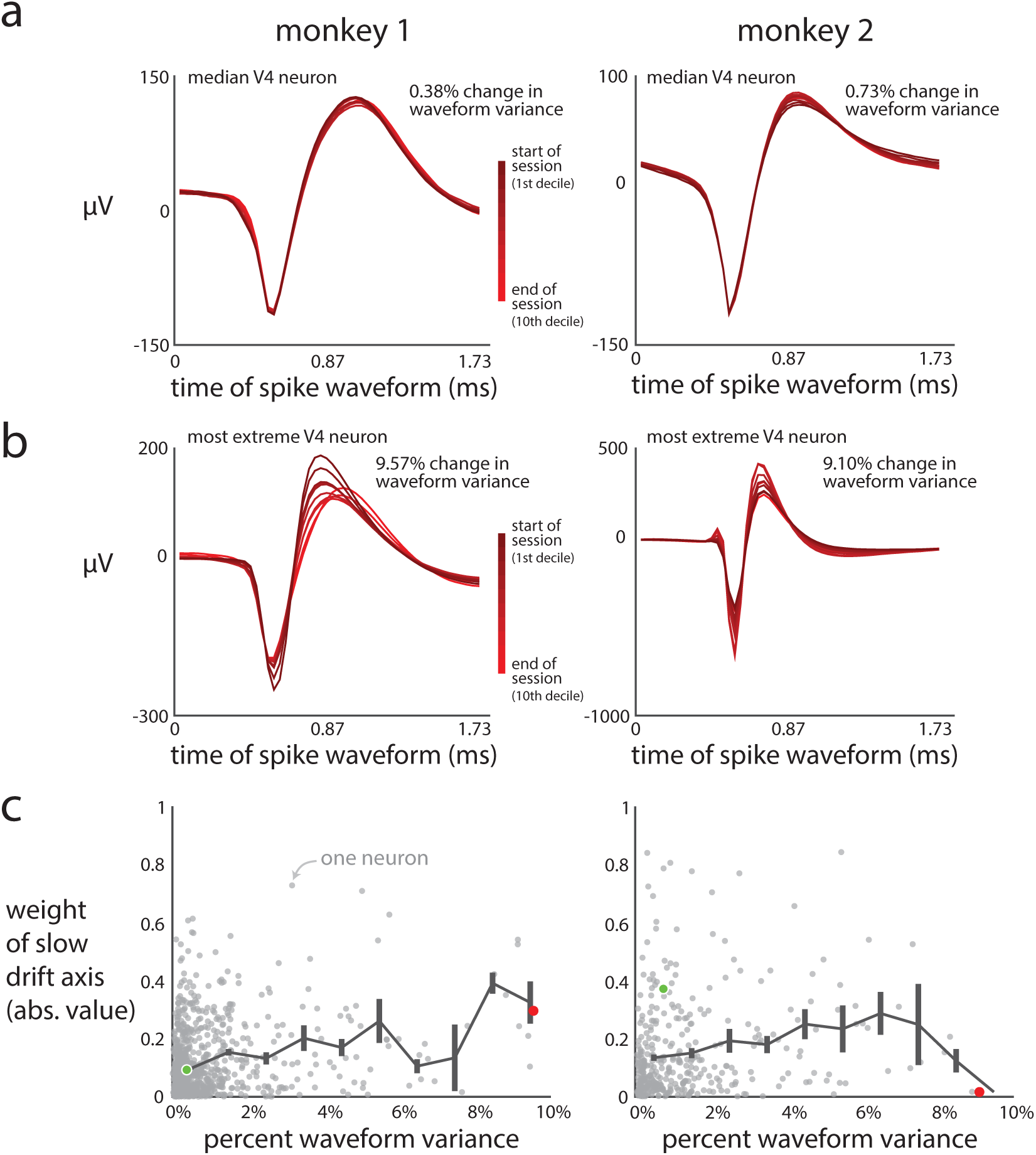
Ensuring the recording stability of the V4 neuron isolations. One potential source of the slow drift could have been recording instability, which might have caused the recorded spike waveforms to gradually change during a session. A changing spike waveform could have affected our spike sorting procedure, leading to a spurious slow drift in neural activity. To ensure this was not the case, we assessed how much a neuron’s spike waveform changed throughout a session with the percent waveform variance metric (see Methods). **a**. Spike waveforms for V4 neurons with the median percent waveform variance for each monkey. Each trace represents the average waveform for one decile of the session. **b**. Same conventions as in panel **a** except for V4 neurons we analyzed with the largest percent waveform variance (neurons with a percent waveform variance greater than 10% were removed from our analyses). **c**. Having removed neurons with percent waveform variance greater than 10%, we performed two analyses to verify that the identified slow drift was not a product of recording instabilities in the remaining neurons. First we asked whether neurons with highly stable waveforms (i.e., percent waveform variance < 1%) contributed to the slow drift. We found many neurons with a very low percent waveform variance but large weight magnitude (i.e., there are many gray dots with < 1% waveform variance and > 0.2 weight magnitude). Thus, even the most stably-recorded neurons contribute substantially to the identified slow drift. Second, we binned the percent waveform variances into 1% bins, and computed the mean weight magnitude across neurons for each bin (black line). This mean value was relatively flat across bins, suggesting neurons at all percent waveform variance levels contributed to the identified slow drift. Black error bars indicate ±1 s.e.m., and the green and red dots denote the median and most extreme neurons, respectively. These results indicate that the identified slow drift in neural activity could not have been caused by neural recording instabilities.

**Supplementary Figure 4:**
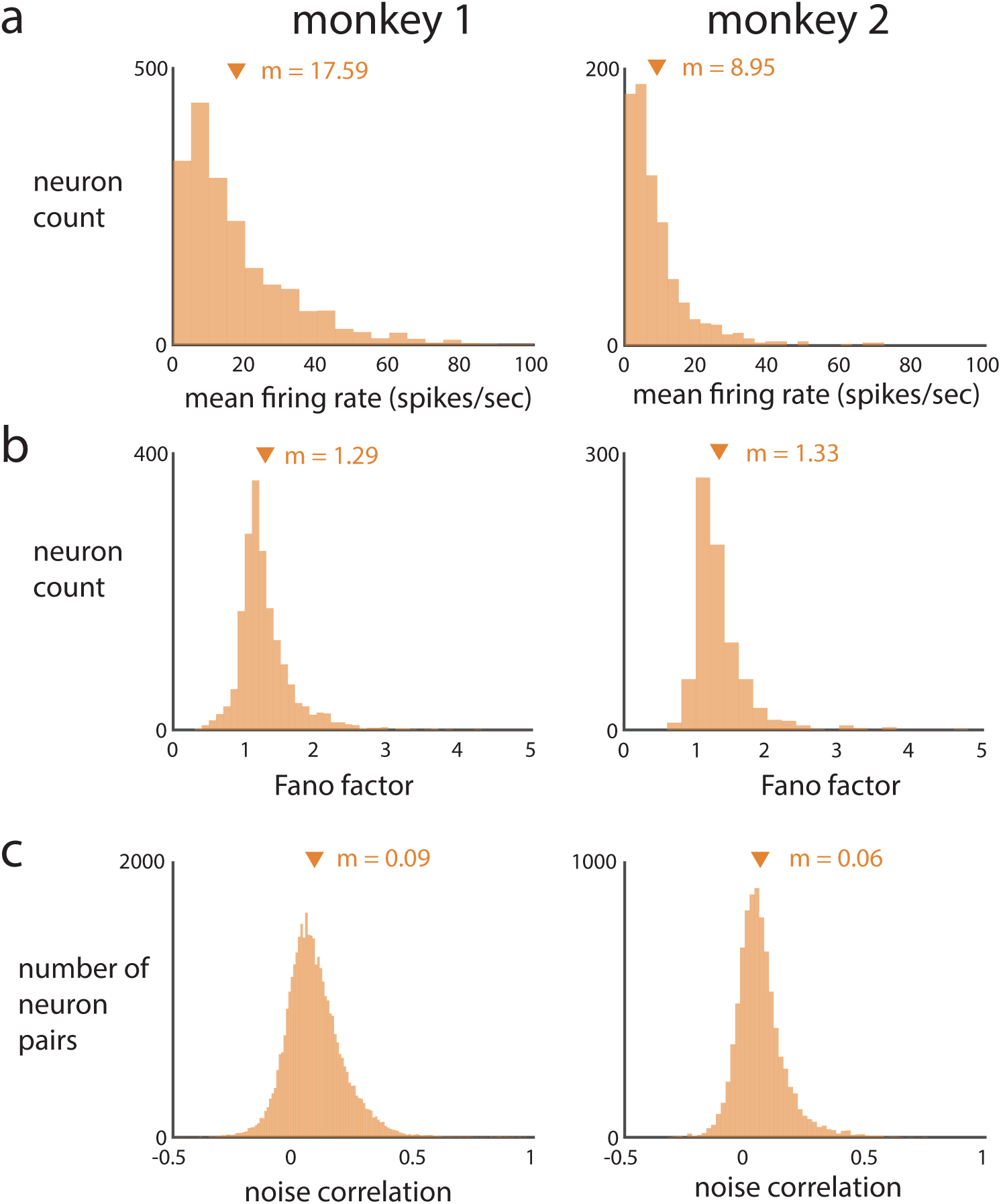
Properties of V4 neural activity were consistent with those observed in previous studies [reviewed in 17]. Spike counts were taken in a 400 ms bin starting 50 ms after stimulus onset. Results for monkey 1 (left column) and monkey 2 (right column) were computed separately for each stimulus orientation (i.e., 45° and 135°) and aggregated across orientations and sessions. **a**. Mean firing rate. **b**. Fano factors (spike count variance divided by mean spike count). **c**. Noise correlations for each pair of simultaneously-recorded neurons. Triangles indicate means (‘m’).

**Supplementary Figure 5:**
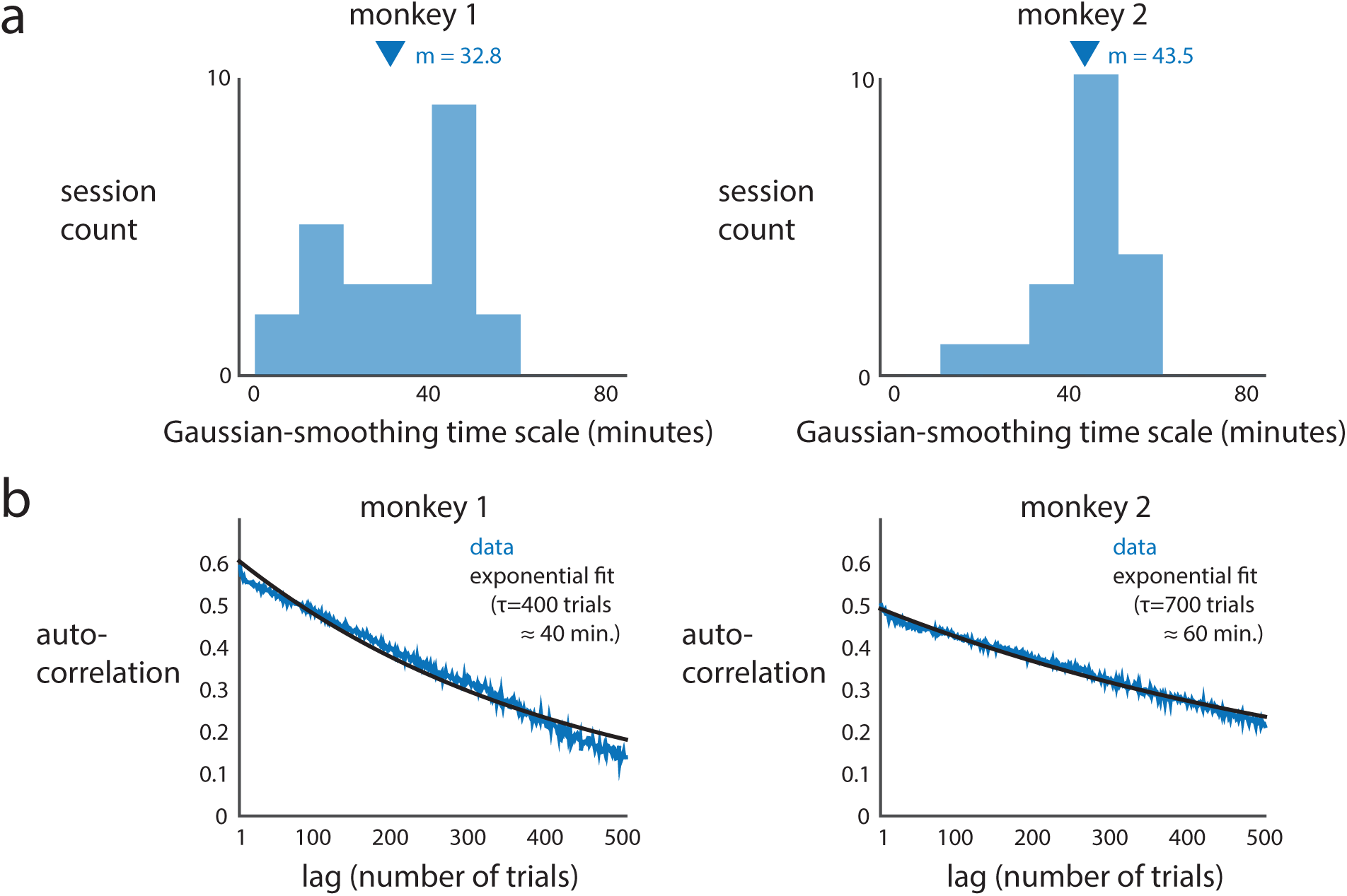
The timescale of the V4 slow drift was approximately 40 minutes. **a**. We estimated the timescale of the V4 slow drift with Gaussian smoothing (same estimation procedure as that for estimating the timescale of hit rate and false alarm rate, see Methods). We found that this timescale for the slow drift of V4 activity (i.e., V4 responses projected onto the slow drift axis) was approximately 40 minutes for each monkey (mean *m* = 32.8, 43.5 minutes for monkeys 1, 2). **b**. We also computed the timescale of the slow drift using a different method. We performed an autocorrelation on the responses projected onto the slow drift axis (blue, averaged over sessions). We fit an exponential function (black) to the autocorrelation curve (blue), and found decay rates of *τ* = 400, 700 trials for monkeys 1, 2. On average, 100 trials took 10.2, 9.4 minutes for monkeys 1, 2, indicating that the timescale of the slow drift was ∼50 minutes (∼40, ∼60 for monkeys 1, 2). One caveat is that the autocorrelation requires that responses have equal spacing between time points, but this was not the case for our data: trials could be separated by different lengths in time. Taken together, the results of the Gaussian smoothing analysis and the autocorrelation analysis suggest that much of the response variation can be explained by a slow drift with a timescale of 40 minutes or greater.

**Supplementary Figure 6:**
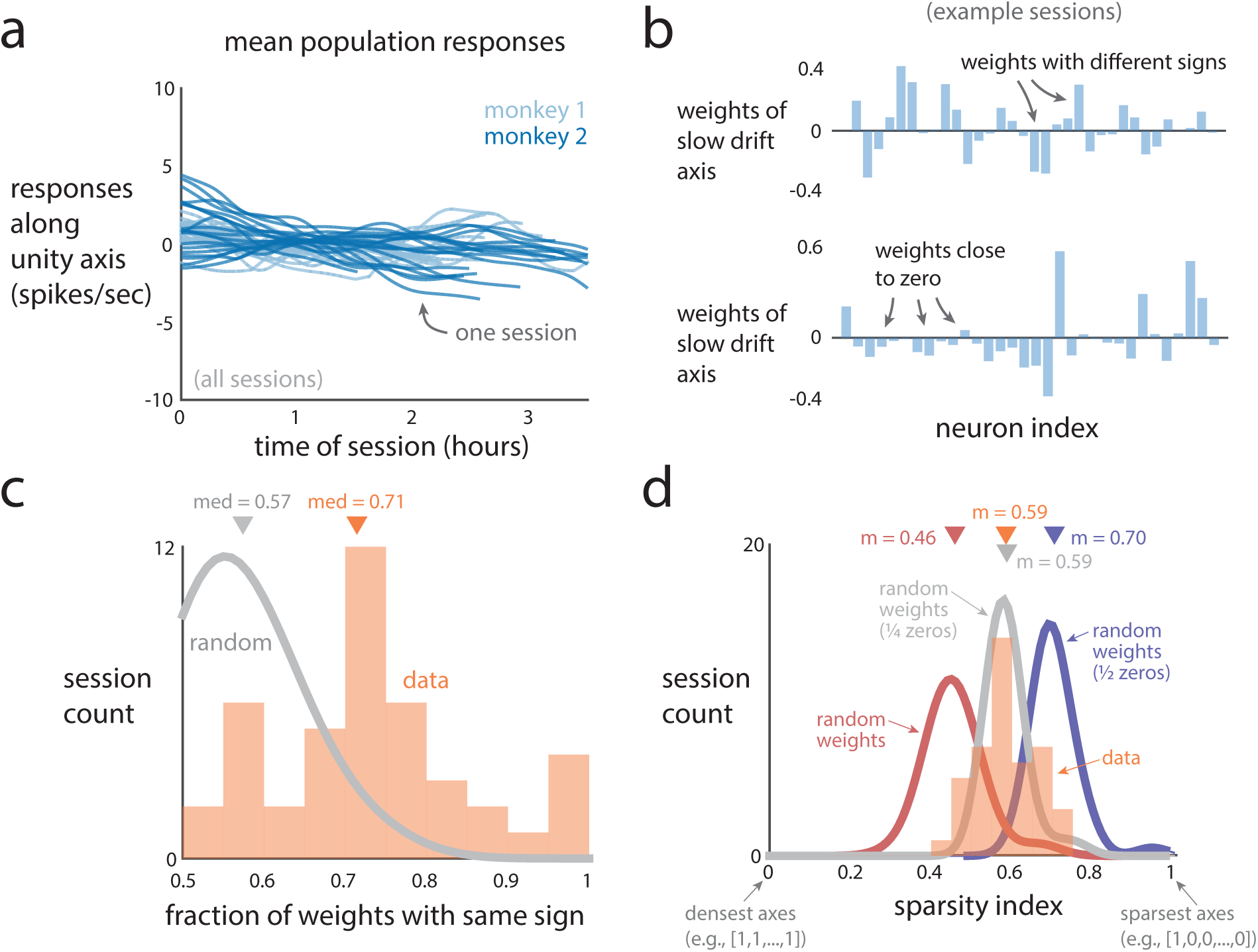
The slow drift’s effects on firing rates may be missed if one simply averages firing rates across neurons. **a**. Time courses of the mean population rate for all sessions. The mean population rate was computed by projecting population activity onto the unity axis 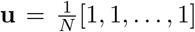 for *N* neurons and then Gaussian smoothed. The slow drift was processed in the same manner except that the slow drift axis was identified with PCA (Fig. 2d). The slow drift varied substantially more than the corresponding mean population rate (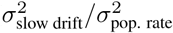 median = 6.9, 2.9 for monkeys 1, 2 and were significantly above 1, *p <* 0.05 for both monkeys, paired permutation test). In addition, the slow drifts were not correlated with their corresponding mean population responses (mean *ρ* = 0.04 ± 0.16, *−*0.04 ± 0.21 for monkeys 1, 2, where ±1 indicates 1 s.e.m.). These results indicate that the slow drift is not apparent when directly averaging the activity across neurons. **b**. Weight vectors of the slow drift axis for two example sessions. In the following two analyses, we asked whether the weights had the same signs and how many weights were close to zero. **c**. We assessed to what extent the slow drifts of neurons were positively or negatively correlated by analyzing the signs of the weights. If some neurons were negatively correlated, it suggests that taking the population average may miss large effects of the slow drift. We found this to be the case, as only 70% of neurons had weights with the same sign. We found this by computing the fraction of weights with the same sign for each session. A fraction of 1 indicates that all neurons have weights with the same sign. A fraction of 0.5 indicates that half the neurons have positive weights, while the other half have negative weights. The median fraction for all sessions was 0.71 (‘data’). For reference, the distribution of fractions for weights whose signs were randomly flipped with equal probability (‘random’, smoothed histogram) had a median of 0.57. **d**. Another important consideration was to what extent the weights of the slow drift axes were sparse (i.e., how many weights had values far from zero). This is important because it could have been the case that only 1 to 2 neurons had large weights and the rest of the weights were close to zero (i.e., high sparsity), implying that the slow drift is not a population effect but rather an effect of a small number of specific neurons. However, this was not the case, as we found that roughly 75% of the recorded neurons slowly drifted by performing the following analysis. We defined a sparsity index for slow drift axis **s** *∈* R*^N^* for *N* neurons as the angle between [*|***s**_1_*|, …, |***s***_N_ |*] (where **s***_i_* is the *i*th element of **s** and *| · |* is the absolute value function) and [1, 1, …, 1] (i.e., the unity axis) divided by the maximally-sparse angle. The maximally-sparse angle is the angle between the [1, 1, …, 1] axis and the [1, 0, …, 0] axis (i.e., a unit axis). A sparsity index of 1 indicates that **s** is highly sparse with many weights that are close to zero. A sparsity index of 0 indicates that **s** is dense (i.e., many weights far from zero). For reference, we computed the sparsity index for randomly-generated vectors where each weight was drawn from a standard Gaussian (red). We computed two other reference distributions whose vectors were generated in the same way as that for the red distribution but a fraction of weights were forced to be zero (gray: 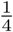 of weights are zero, blue: 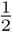 of weights are zero). The distribution of sparsity indices for the slow drift axes (orange, ‘data’) overlapped the most with the 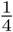-weights distribution (gray), indicating that the activity of roughly 75% of the recorded neurons slowly drifted. Taking **c** and **d** together, we conclude that ∼50% of neurons increase or decrease their activity together, ∼25% either increase or decrease their activity in the opposite manner of the first 50%, and ∼25% have little to no drift.

**Supplementary Figure 7:**
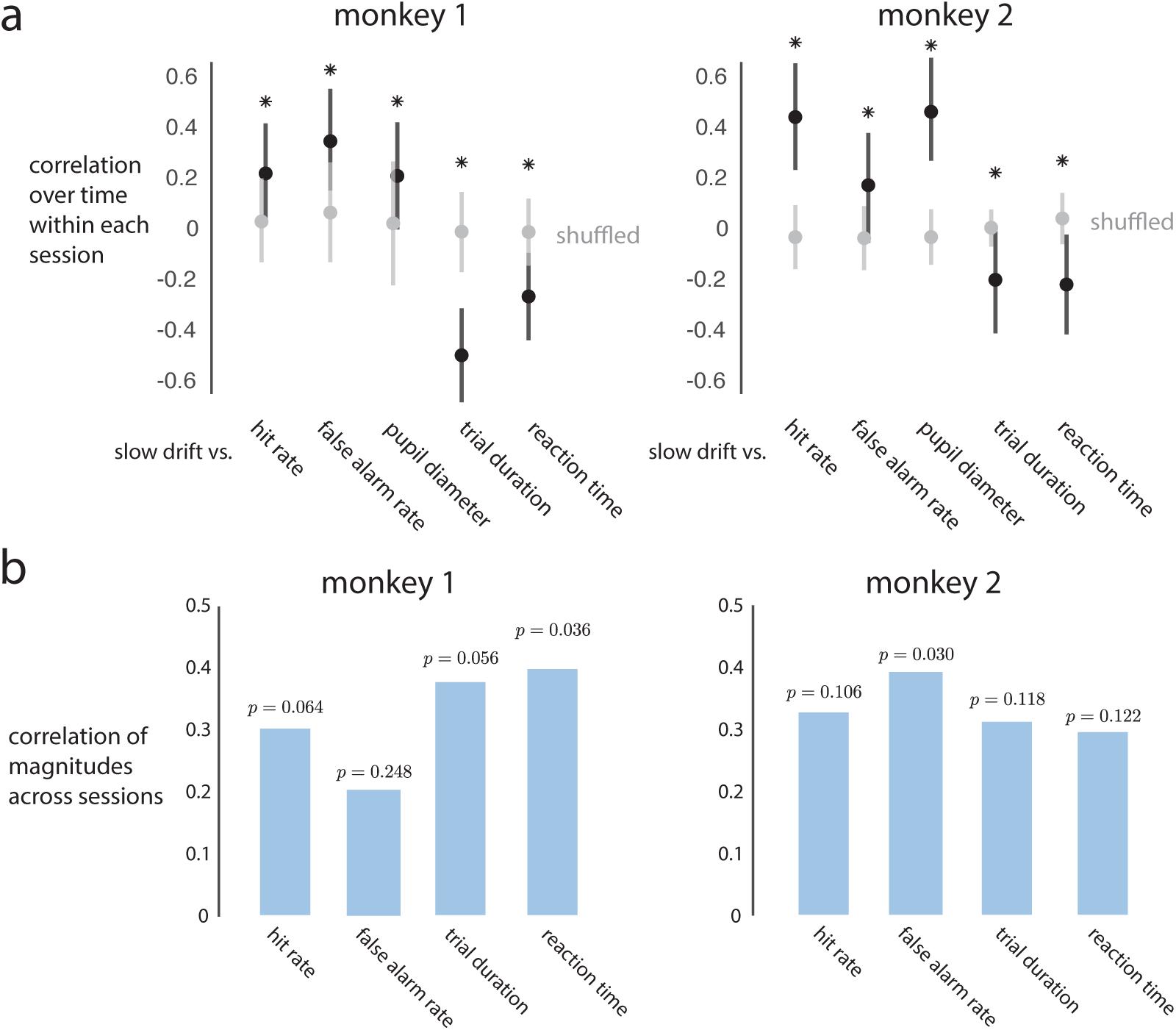
The slow drift covaries with slow fluctuations in behavior. Similar trends for individual monkeys as aggregated results (Fig. 3b and c). Same plotting conventions as in Figure 3b and **c**. **a**. Correlations over time within each session between the V4 slow drift and behavioral variables for each monkey. Correlations match those of aggregated results (Fig. 3b). Asterisk denotes significance over chance levels (*p <* 0.05, permutation test). **b**. Correlations between magnitude of slow drift and magnitudes of behavioral variables for each monkey. All correlations are positive and close to significant (*p*-values are shown, permutation test).

**Supplementary Figure 8:**
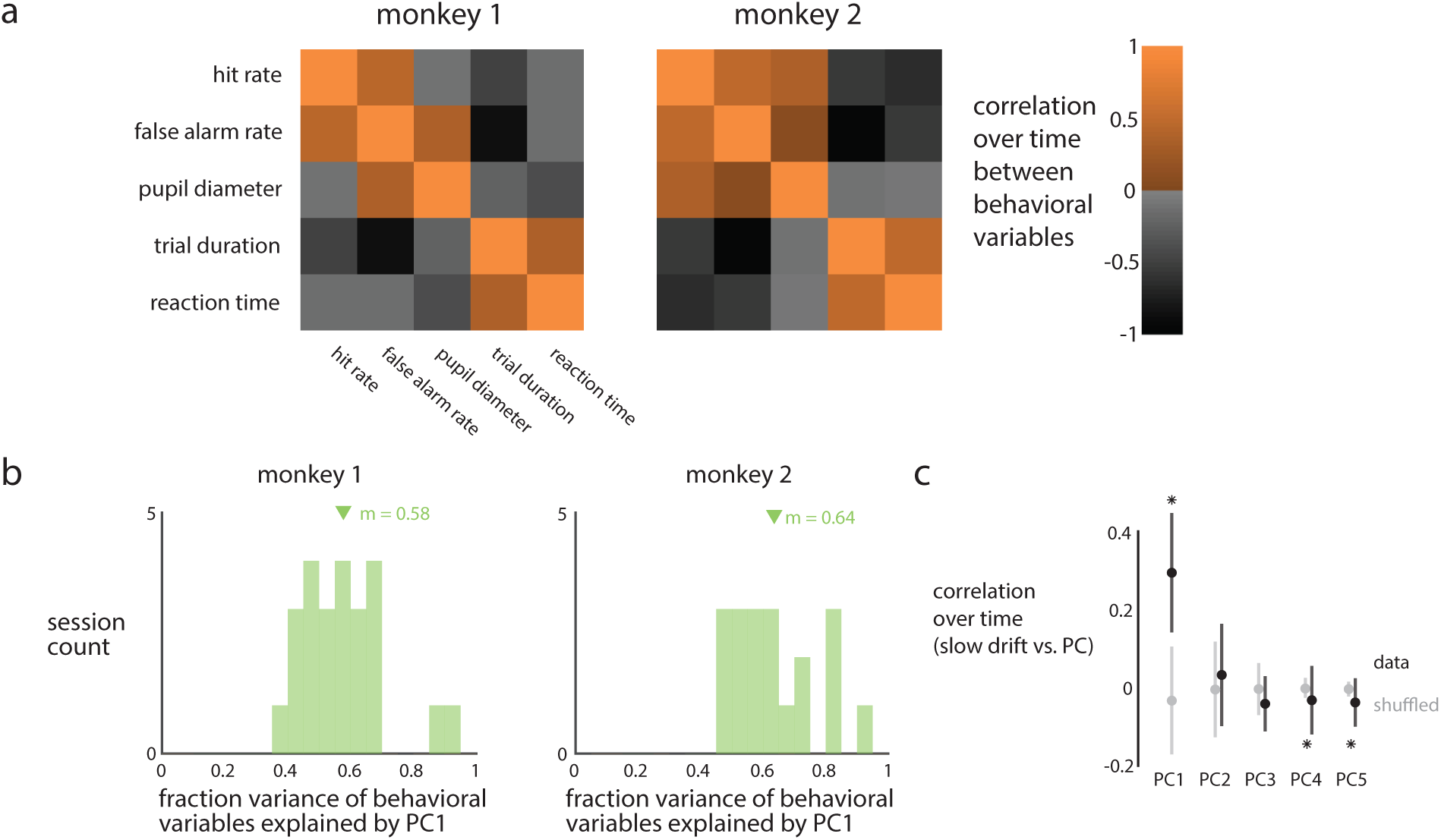
Behavioral variables primarily covaried along one dominant pattern, and the slow drift was most correlated with this pattern. **a**. We investigated the correlations over the time within each session between the five behavioral measures for each monkey. We found a similar trend as that of the correlations between the slow drift and behavioral variables (Fig. 3b). Namely, hit rate, false alarm rate, and pupil diameter were mostly positively correlated; trial duration and reaction time were positively correlated; and these two groups were negatively correlated. **b**. We next asked to what extent the covariance matrices in **a** could be explained by one co-fluctuation pattern among the behavioral variables. We applied PCA to the behavior variables, and found that the top principal component (PC1) explained 58%, 64% of the variance of the behavioral variables for monkeys 1, 2. This indicates that there is a dominant co-fluctuation pattern among the behavioral variables. We confirmed that the weights of this dominant pattern matched the pattern of correlations in **a** (i.e., the weights were close to [1, 1, 1*, −*1*, −*1], which matched the groupings of correlations). **c**. Finally, we asked to what extent did the slow drift covary with different co-fluctuation patterns of the behavioral variables. The slow drift covaried most strongly with the first principal component (PC1), significantly greater than that of shuffled data (*p <* 0.002, permutation test). This relationship was weaker for the other PCs (PC2-PC5), and in the opposite direction for the weakest PCs (PC4 and PC5, asterisk corresponds to *p <* 0.05, permutation test). Black and gray dots indicate median correlations across sessions. Error bars indicate bootstrapped 90% confidence intervals. These results indicate that a single dominant pattern explains the co-fluctuations of the behavioral variables, and that the slow drift is most correlated with this pattern.

**Supplementary Figure 9:**
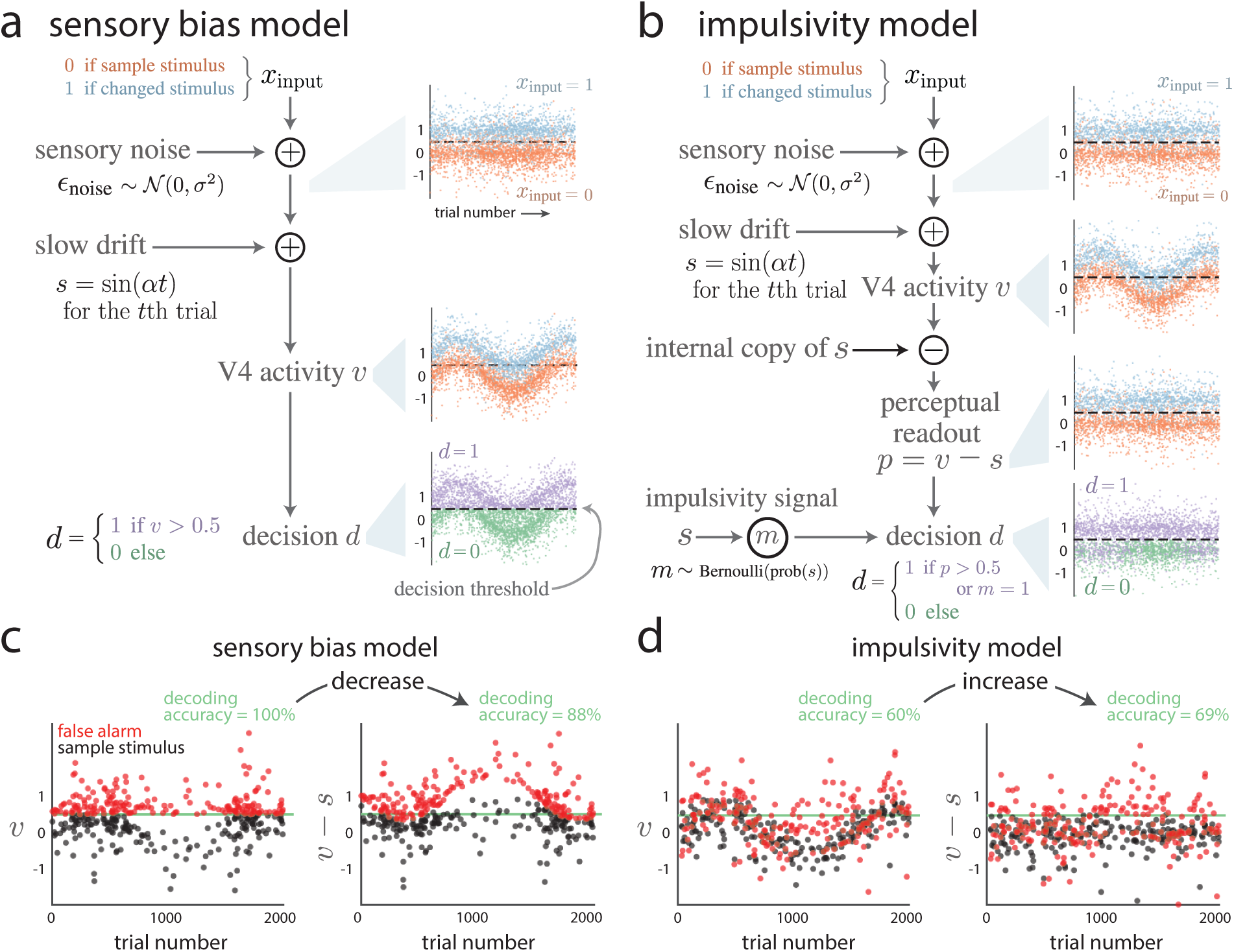
Simulation procedure of the sensory bias model and the impulsivity model, and illustration of the decoding analysis in Figure 6e and **j**. Simulations have the same trial structure as that of the orientation-change detection task. Each trial consists of a sequence of stimulus inputs *x*_input_ *∈ {*0, 1*}*, where *x*_input_ = 0 indicates no stimulus change and *x*_input_ = 1 indicates a stimulus change. The trial sequence starts with *x*_input_ = 0 and has a 40% chance of changing from *x*_input_ = 0 to *x*_input_ = 1 for each consecutive input. The correct output of the model is to indicate decision *d* = 0 if *x*_input_ = 0 and *d* = 1 if *x*_input_ = 1. **a**. Simulation procedure of the sensory bias model. Sensory noise *∊*_noise_ is added to *x*_input_ (top right panel depicts *x*_input_ + *∊*_noise_). Slow drift *s* is then added to form the simulated V4 activity *v* = *x*_input_ + *∊*_noise_ + *s* (middle right panel depicts *v*). Finally, the decision *d ∈ {*0, 1*}* is based on *v* (bottom right panel depicts *v* but colored based on decision *d*). Dashed black lines indicate the decision threshold used to determine decision *d*. For the sensory bias model, any V4 activity *v* above the decision threshold of 0.5 leads to a “saccade” (i.e., *d* = 1). **b**. Simulation procedure of the impulsivity model. V4 activity is simulated in the same manner as that for the sensory bias model (i.e., top right two panels are the same in **a** and **b**). An internal copy of *s* is removed from the readout of the V4 activity to form the perceptual readout *p* = *v − s* (right panel, third from the top, depicts *p*). Decision *d* is based on the perceptual readout *p* passing a decision threshold (i.e., *d* = 1 if *p >* 0.5) or an impulsivity signal *m* (i.e., *d* = 1 if *m* = 1). It is more likely for impulsivity signal *m* to equal 1 if slow drift *s* has a higher value. An important point is that impulsivity signal *m* influences decision *d* independent of perceptual readout *p*, leading to some decisions *d* = 1 that are not caused by the perceptual readout *p* passing the decision threshold (bottom right panel, some purple dots are below the dashed line, cf. bottom right panel in **a**). **c**. Illustration of the analysis procedure for decoding false alarms from the simulated V4 activity for the sensory bias model. We consider only false alarm trials, and compare simulated V4 activity *v* corresponding to the last stimulus input of the trial (i.e., when a false alarm occurred because *d* = 1 when *x*_input_ = 0, red dots) versus the preceding stimulus input (i.e., a correctly rejected stimulus input because *d* = 0 when *x*_input_ = 0, black dots). The red dots form a subset of the purple dots in the bottom right panel in **a**, and the black dots form a subset of the green dots. We decode V4 activity *v* (i.e., with the slow drift) and *v − s* (i.e., with the slow drift subtracted) using a threshold decoder (green lines, linear SVM decoder). For the sensory bias model, subtracting the slow drift *decreases* the decoding accuracy (i.e., decoding accuracy is higher when decoding *v* than when decoding *v−s*). This is because the slow drift biases the sensory evidence *v* to be closer or further from the decision threshold (**a**, bottom right panel), and removing this bias discards information that V4 activity has about the decision. **d**. Same analysis procedure as in **c**, except for the impulsivity model. Subtracting the slow drift *increases* the decoding accuracy (i.e., decoding accuracy is lower when decoding *v* than when decoding *v − s*). This is because the slow drift is removed from perceptual readout *p* (**b**, right panel, third from top). Still, the slow drift influences many decisions unrelated to the perceptual readout (e.g., many false alarms occur even when *p* is below the decision threshold because *m* = 1). Here, the ability of *v* to predict the occurrence of a false alarm within a trial comes from the dependence of *d* on *p >* 0.5 and not from the dependence of *d* on *m* = 1. This is because we only consider a false alarm and its preceding correctly-rejected stimulus flash within a trial (i.e., the slow drift is held constant within a trial), and not a false alarm and a correctly-rejected stimulus flash from any trial (for which the slow drift may vary). Note that the decoding accuracies here are higher than those in Figure 6 because here we did not include decision output noise (see Methods) in order to better illustrate the differences between the two models. Including this output noise (which accounts for the fact that we recorded only a small fraction of the neurons in the decision-making circuit, and other unobserved sources of noise are likely) yields overall decoding accuracies of the models (Fig. 6**e** and **j**) more similar to those of the real data (Fig. 6**f**).

**Supplementary Figure 10:**
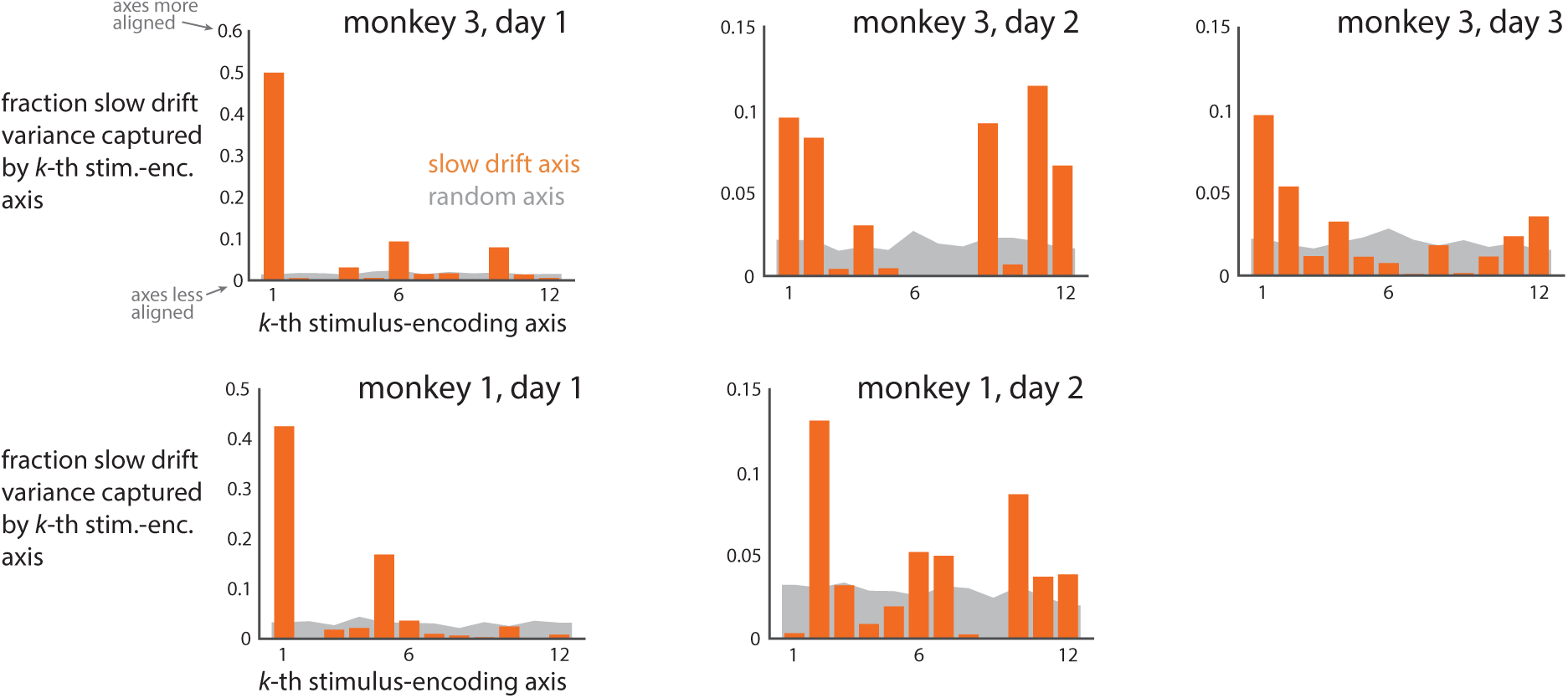
The slow drift axis overlapped with the top stimulus-encoding axes. Same convention as Figure 7**d**, shown for each of the five sessions individually. We asked whether the slow drift axis overlapped with any of the stimulus-encoding axes (i.e., at least one orange bar above gray) versus no overlap (i.e., no orange bar above gray). We computed the fraction of the slow drift variance captured by the top 12 stimulus-encoding axes (variance was summed across the top 12 axes), and found that these top 12 stimulus-encoding axes more closely overlapped with the slow drift axis than with a randomly-oriented axis for each session (*p <* 0.002, proportion of random runs with fractions above the fraction of the slow drift). The sum of these fractions (orange bars) equal the fractions reported in Fig. 7**e**. We chose 12 because this number of stimulus-encoding axes captured a large fraction of stimulus response variance (i.e., variance of repeat-averaged responses taken across images) for each session (62%, 60%, 59% for days 1, 2, 3 of monkey 3, and 70%, 72% for days 1, 2 of monkey 1). These results indicate that the slow drift tends to lie along stimulus-encoding axes, and thus the slow drift could corrupt the fidelity to which V4 activity encodes relevant image statistics relevant to downstream areas. This potential corruption of sensory encoding motivates the removal of the slow drift in order for downstream readout areas to better recover the relevant stimulus information.

**Supplementary Figure 11:**
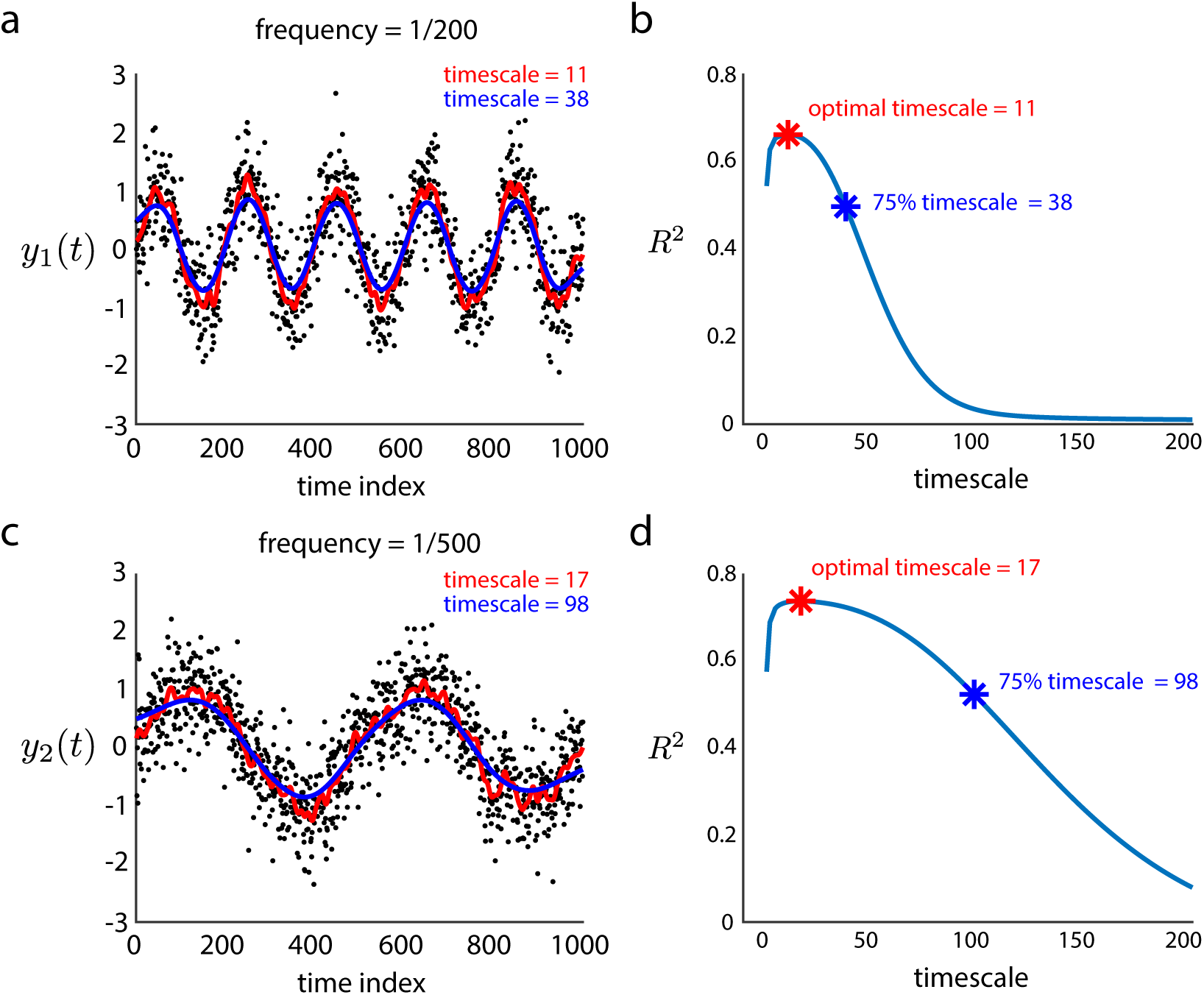
Intuition of our approach for estimating the timescale of hit rate, false alarm rate, and the slow drift. **a**. We simulate a signal 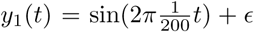, where *t* = 0, …, 1000 represents time (e.g., in minutes) and *∊ ∼ N* (0*, σ*) for *σ* = 0.5 is the noise that is independent at each time point. We simulated 1, 000 time points of this signal (black dots). To estimate this signal’s timescale, we used Gaussian smoothing and considered different standard deviations as candidate timescales (red and blue lines denote smoothed estimates with different timescales). We chose this approach over a Fourier analysis, because this approach allows for the unequal spacing between time points, which occurred in the neural data (i.e., trials were initiated by the animal and inter-trial periods need not have the same durations). **b**. For each candidate timescale, we computed the cross-validated performance *R*^2^ by predicting each held-out data point by a Gaussian weighted average of its neighboring data points (held-out data were chosen randomly for 10 fold cross-validation). We found that the candidate timescale that achieved the largest *R*^2^ (the optimal timescale, red asterisk) was small, and that a much longer timescale still achieved 75% of the largest *R*^2^ (blue asterisk). This suggests that although a small candidate timescale (in this case, 11) may be optimal for Gaussian smoothing (because it can capture both low and high frequency components), it does not necessarily reflect the timescale of the most dominant component (e.g., a low frequency of 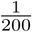). Instead, the timescale of this dominant, low-frequency component is reflected by how slowly the *R*^2^ curve drops off as the candidate timescale increases beyond the optimal timescale. Indeed, the 75% timescale of 38 better reflects the low frequency of 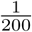. **c**. To confirm this intuition, we simulated *y*_2_(*t*) in the same way as *y*_1_(*t*) except with a lower frequency of 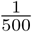. We expected a slower drop off in *R*^2^ for *y*_2_(*t*) than *y*_1_(*t*). **d**. We found that the optimal timescale (red asterisk) for *y*_2_(*t*) was not representative of the dominant frequency of 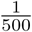. However, the 75% timescale of 98 did reflect this low frequency, indicating the drop in *R*^2^ was slower for a signal with a lower frequency (compare blue curves in **b** and **d**), as expected. Thus, we choose to report the candidate timescale that achieves 75% of the largest *R*^2^ for two reasons. First, this timescale appears to be a more interpretable estimate of the timescale of the most dominant component of the signal (e.g., a timescale of 98 better represents *y*_2_(*t*) in **c** than a timescale of 17). Second, this timescale is more likely to avoid the preference of Gaussian smoothing to choose smaller candidate timescales as the optimal. We only used this approach to report the timescales in the main text for hit rate, false alarm rate, and the slow drift; the estimated timescales do not affect results presented in any figure.

**Supplementary Figure 12:**
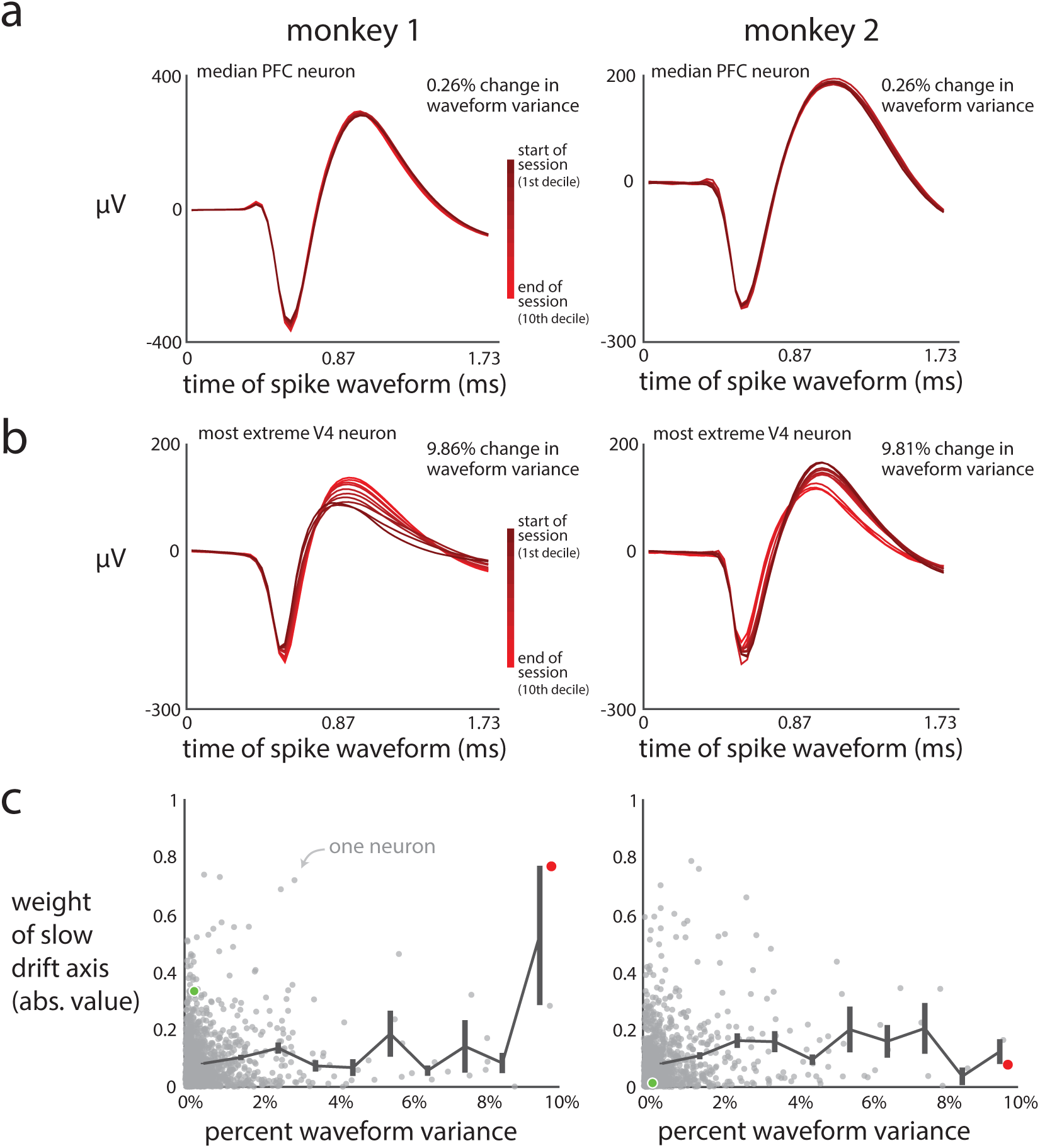
Ensuring the recording stability of the PFC neuron isolations. PFC neurons were subjected to the same neuron removal criteria (*<* 10% waveform variance) as were the V4 neurons. All conventions were the same as in Supplemental Figure 3. These results indicate that any slow drift observed in the activity of the analyzed PFC neurons was not caused by neural recording instabilities.

**Supplementary Figure 13:**
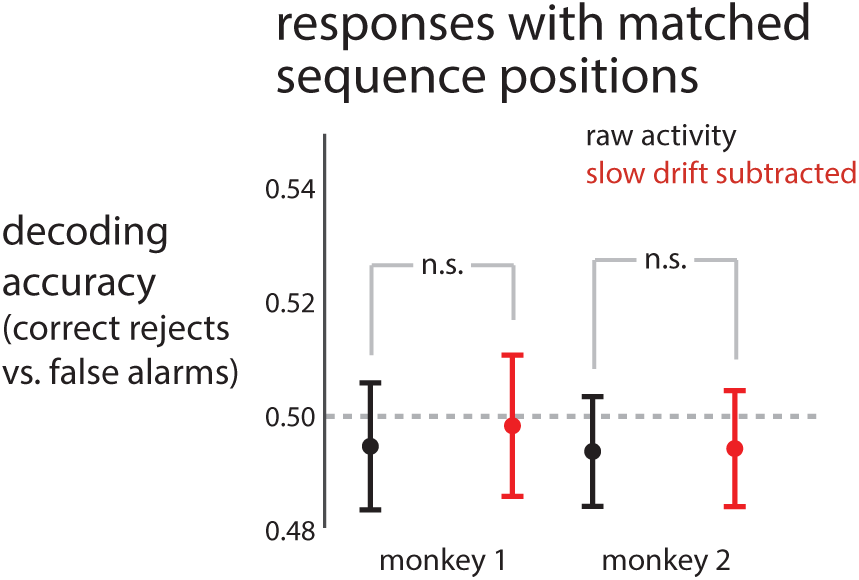
Controlling for visual response adaptation when decoding false alarms versus correct rejects from V4 responses. It could be the case that adaptation-like effects [48] led to an increase in our decoding accuracy in Figure 6**f**, rather than the false alarm itself. To control for this, for each false alarm trial, we identified a nearby matching trial (which need not be a false alarm trial) whose sequence length *M* was at least one flash longer than the sequence length *K* of the corresponding false alarm trial (i.e., *M > K*). We then decoded responses to the two stimulus flashes in positions *K −* 1 and *K* from the matched trial. Because these matched responses come from two correctly-rejected stimulus flashes, we expect our ability to decode these responses to be at chance, unless adaptation-like effects are at play. Indeed, when we decoded these matched responses, we found decoding accuracy close to chance (dots close to gray dashed line). In addition, we found we found no difference in decoding accuracy (*p* = 0.572, *p* = 0.745 for monkeys 1, 2, paired permutation test) when comparing decoding accuracy using the raw activity and the slow drift-subtracted activity, unlike the increase in decoding accuracy observed in responses from false alarm trials (Fig. 6**f**). Dots indicate means across sessions, and error bars indicate ±1 s.e.m. These results indicate that the results in Figure 6**f** are not due to adaptation-like effects.

